# Intra-Nuclear Tensile Strain Mediates Reorganization of Epigenetically Marked Chromatin During Cardiac Development and Disease

**DOI:** 10.1101/455600

**Authors:** Benjamin Seelbinder, Soham Ghosh, Alycia G. Berman, Stephanie E. Schneider, Craig J. Goergen, Sarah Calve, Corey P. Neu

## Abstract

Environmental mechanical cues are critical to guide cell fate. Forces transmit to the nucleus through the Linker of Nucleo- and Cytoskeleton (LINC) complex and are thought to influence the organization of chromatin that is related to cell differentiation; however, the underlying mechanisms are unclear. Here, we investigated chromatin reorganization during murine cardiac development and found that cardiomyocytes establish a distinct architecture characterized by relocation of H3K9me3-modified chromatin from the nuclear interior to the periphery and co-localization to myofibrils. This effect was abrogated in stiff environments that inhibited cardiomyocyte contractility, or after LINC complex disruption, and resulted in the relocation of H3K27me3-modified chromatin instead. By generating high-resolution intra-nuclear strain maps during cardiomyocyte contraction, we discovered that the reorganization of H3K9me3-marked chromatin is influenced by tensile, but not compressive, nuclear strains. Our findings highlight a new role for nuclear mechanosensation in guiding cell fate through chromatin reorganization in response to environmental cues.

## INTRODUCTION

Mechanics of cell microenvironments play an important role in directing cell differentiation during development^1^ and maintaining tissue health during adulthood^2^. Changes in mechanical properties due to acute trauma, chronic conditions, or genetic predispositions lead to cellular degeneration and result in a range of pathologies^3^, including cardiac hypertrophy^4,5^. Further, regenerative medicine aims to engineer suitable microenvironments to guide cell fates for enhanced tissue regeneration. However, little is known about the underlying mechanisms that facilitate cell differentiation in response to environmental cues.

The nucleus is thought to be an essential mechanosensitive organelle^6–10^ as it is tightly connected to all parts of the cytoskeleton through LINC (Linker of Nucleo- and Cytoskeleton) complexes^11^ comprised of proteins that span the inner (SUNs) and outer (nesprins) nuclear membranes^12,13^. Studies on isolated nuclei demonstrated that the nucleus alone can respond to stretch; however, only when engaged via LINC complexes^14^. Mutations in LINC complex and nuclear envelope proteins are also related to developmental disorders^15^, particularly in mechanically active tissues such as cardiac and skeletal muscle^16,17^. In addition, there is a direct relation between nuclear architecture and cell differentiation. Chromatin organization changes from an unstructured organization in the zygote to a cell type-specific organization during development^18–22^. The 4D Nucleome Project aims to generate spatial maps of human and mouse genomes to better understand this relationship^23^. Since the nucleus makes up a large portion of the cell, type-specific nuclear morphology can also have direct implications for cellular functions as has been described for plasma cells^24^, neutrophil granulocytes^25^, T-cells^26^ and photoreceptor cells^27^. Together, this suggests that biophysical signals from the cell environment might guide cell behavior through spatial rearangment of chromatin; however, no study has investigated the effect of nuclear strains on chromatin organization.

To bridge this gap, we investigated the reorganization of chromatin during cardiomyocyte (CM) development and pathology in mice. CMs show poor contractility and inhibited differentiation on substrates that are stiffer than their native environment^28–30^. Due to their well-investigated behavior in response to substrate stiffness and contraction-mediated deformation of nuclei, CMs provided a good model to investigate the relation between micro-environment, cell differentiation, and chromatin organization, as well as the potential role of nuclear mechanosensation in these processes. We documented the establishment of a distinct nuclear architecture during development and found evidence that tensile nuclear strains, transferred from myofibrils via LINC complexes, guided the reorganization of H3K9me3-modified chromatin to establish this architecture. Reduction of nuclear strains in stiff environments or disruption of LINC complexes inhibited the formation of the CM nuclear architecture and lead to the rearrangement of H3K27me3-modified chromatin instead. To find direct evidence for a link between nuclear deformation and chromatin reorganization in CMs, we used a recently developed method called *deformation microscopy*^31,32^ to generate high resolution intra-nuclear strain maps from microscopy-based image series recorded during CM contractions and devised a workflow to map these strains back to chromatin regions with different epigenetic modifications. Overall our findings suggest a new role for nuclear mechanosensation in CMs in which the nucleus integrates mechanical signals from the environment through the reorganization of epigenetically marked chromatin to guide and stabilize cell differentiation.

## RESULTS

### CMs Adopt a Distinct Nuclear Architecture During Development

To study the relationships between functional microenvironments, nuclear morphology, and chromatin organization, we used adult H2b-eGFP mice to analyze nuclear architectures in primary cells of tissues with different stiffness properties and that undergo a broad range of mechanical challenges (**Fig. 1a**)^33^. While all cell types showed distinct nuclear architectures, nuclei of CMs had an elongated morphology with chromatin accumulated at the nuclear periphery and inner cavities that appeared almost void of chromatin. In contrast, cardiofibroblasts (CF), which shared a similar mechanical environment and had an elongated nuclear morphology, showed a homogeneous distribution of chromatin throughout the nucleus. Analysis of late stage (E)18.5 embryonic hearts revealed that the nuclear architecture observed in adult CMs was not present in embryonic CMs (**Fig. 1b**), suggesting that the adult nuclear phenotype forms postnatally when a sudden increase in cardiac activity triggers CM maturation.

**Fig. 1.**
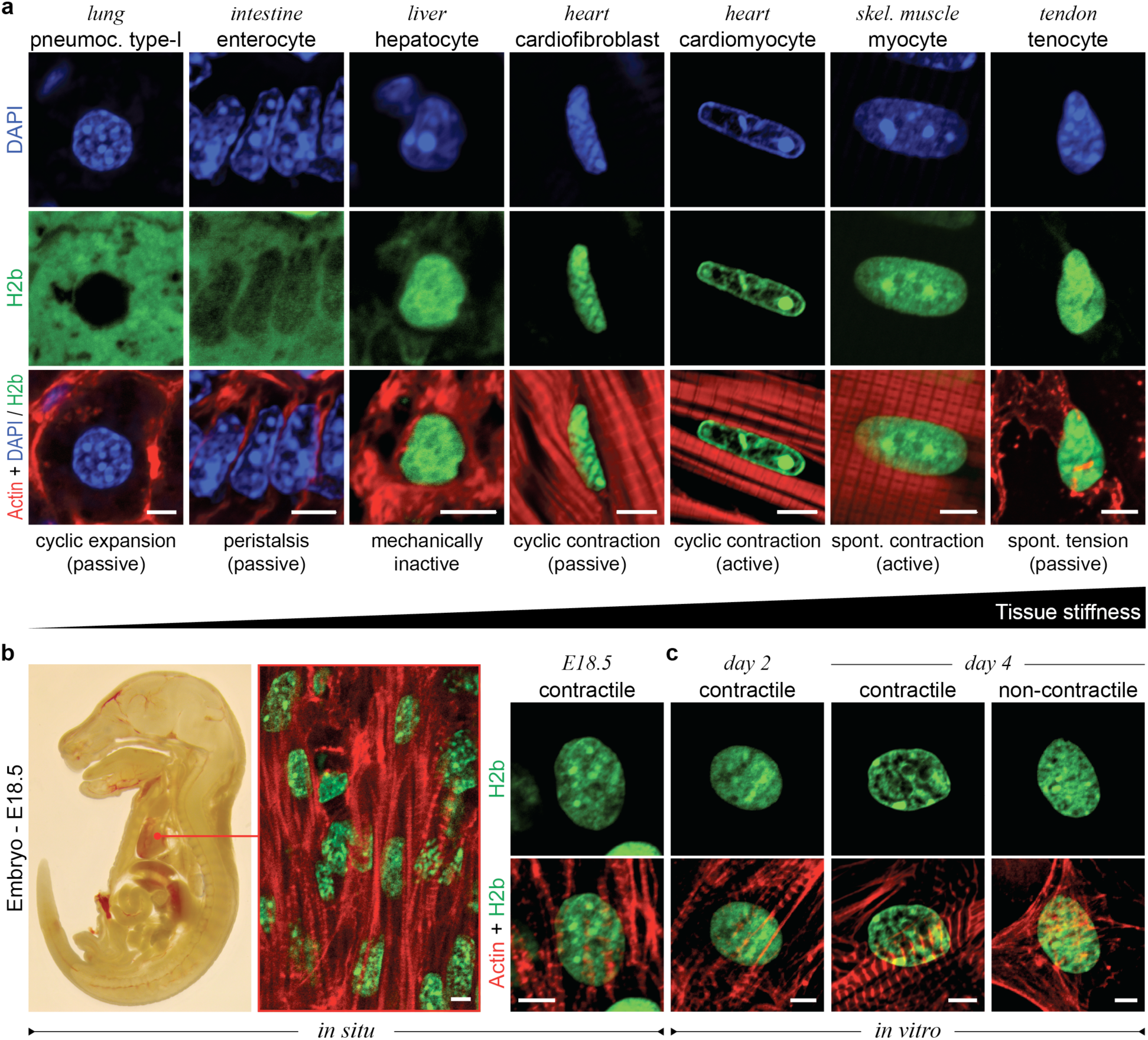
Cardiomyocytes adopt a distinct nuclear architecture with high amounts of peripheral chromatin during development. **a)** Tissues with diverse mechanical characteristics were harvested from adult H2b-eGFP mice and stained for actin. DAPI was used as DNA counterstain for soft tissues with weak H2b-eGFP fluorescence. Adult CMs had an elongated nucleus with a high ratio of peripheral chromatin compared to other cell types. **b)** Embryos from H2b-eGFP mice were harvested at embryonic day (E)18.5, sectioned and stained for actin. Left: whole embryo mid-section. Middle: close-up of embryonic cardiac tissue from mid-section. Right: close-up of an embryonic CM nucleus, which showed a diffuse nuclear organization unlike adult CMs. **c)** Embryonic cardiac cells were isolated from (E)18.5 H2b-eGFP mice hearts and plated on soft (13 kPa) PDMS substrates for two or four days. Embryonic CMs with contractile myofibrils showed a change in nuclear organization at day four. All scales=5 µm.

To analyze the formation of the adult nuclear phenotype in CMs during development, we isolated embryonic cardiac cells from (E)18.5 mouse hearts by using an optimized mixture of ECM-specific peptidases (see methods for details) to achieve high cell yields and high viability compared to existing tryptic methods^34^. Isolated cells were plated on soft (13 kPa) silicone substrates (polydimethylsiloxane, PDMS), coated with basement membrane proteins, to mimic the mechanical environment of adult hearts^28,29,35^. The resulting cardiac co-culture contained a high percentage (61%) of embryonic CMs, assessed by the formation of myofibrils, even without enrichment of CMs through pre-plating (**Extended Fig. 1** and **Fig. 2a**). Embryonic CMs grew in connected clusters, started coordinated contractions within 24 hours of plating, and remained at a high ratio over a four-day culture period (**Fig. 2a**). After two days in culture, nuclear phenotypes of CMs still appeared diffuse with no distinctive accumulation of heterochromatin at the nuclear envelope (**Fig. 1c**). However, after four days, embryonic CMs exhibited intranuclear cavities devoid of chromatin and an overall shift of chromatin towards the nuclear periphery like adult CMs (**Fig. 1a**). Interestingly, non-contractile CFs present in the culture continued to display a diffuse nuclear architecture more similar to adult CFs. These findings provided further support for a link between chromatin organization and cell differentiation and suggest that CMs form a cell type-specific nuclear architecture during development.

**Fig. 2.**
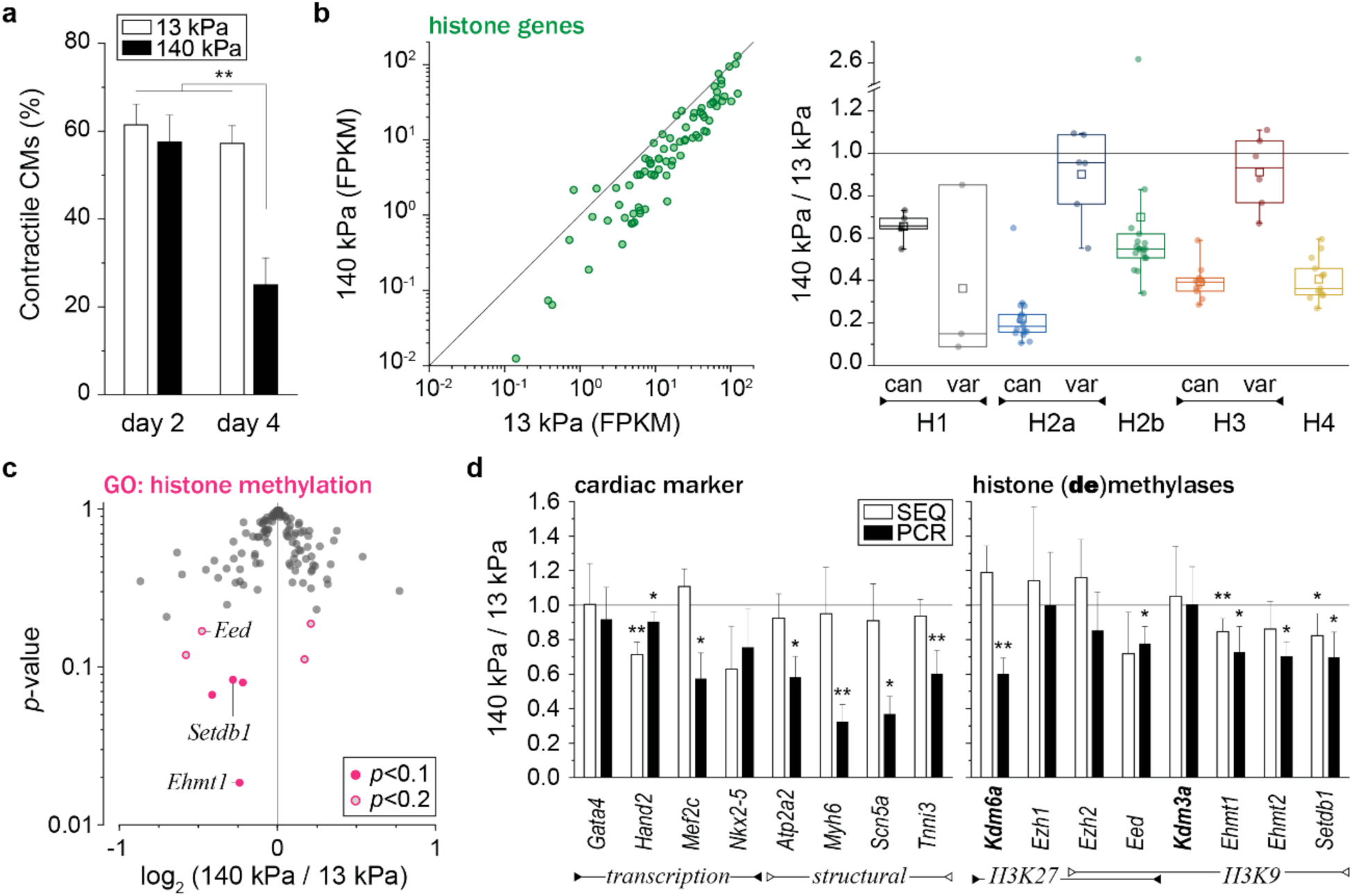
Substrate stiffness affects gene expression of histones and histone modifying enzymes in embryonic cardiac cultures. Cells were isolated from (E)18.5 H2b-eGFP embryo hearts and cultured on soft (13 kPa) or stiff (140 kPa) PDMS. **a)** After two and four days, cultures were stained for actin (see Extended Fig.1) and the ratio of CMs with contractile myofibrils to non-contractile CFs was determined. The percentage of contractile CMs was significantly reduced after four days on stiff substrates compared to soft; SEM, n=9 from 3 exp., 2W-ANOVA: ** p<0.01. **b)** Total RNA was collected after four days of culture. RNAseq analysis revealed that most of the 82 expressed histone genes were downregulated on stiff PDMS. Canonical histones (can) were consistently downregulated for all histone families while non-canonical histone variants (var) showed varying levels of expression changes between substrates; n=4; FPKM: Fragments Per Kilobase of transcript per Million mapped reads. **c)** Volcano plot of genes associated with the gene ontology term *histone methylation* (GO:0016571) as determined by RNAseq. Indicated are genes coding for H3K9 methylases, which were amongst the most significantly altered (see also Extended Table 2). **d)** PCR validation of RNAseq (SEQ) data verified downregulation of H3K9 methylating genes and showed that cardiac transcription and structural marker were decreased on stiff substrates. H3K9 demethylase *Kdm3a* and H3K27-specific methylase *Ezh1* showed no change while H3K27 demethylase *Kdm6a* was downregulated, indicating inverse methylation activity between H3K9 and H3K27 for cardiac cells on stiff substrates; SD; n=4; T-test: * p<0.05, ** p<0.01.

### Substrate Stiffness Affects Histone and Epigenetic Enzyme Expression in Embryonic Cardiac Cells In Vitro

To better understand the relationship between CM-specific nuclear architecture and CM differentiation, we next screened for changes in gene expression related to chromatin remodeling in an *in vitro* model of cardiac dedifferentiation. Embryonic CMs show reduced contractility and dedifferentiate in environments that are stiffer than native cardiac tissue (12 ± 4 kPa)^28–30^. We verified these results by analyzing the ratio of contractile CMs to non-contractile CFs, assessed by the formation of contractile myofibrils, of embryonic cardiac cells plated on stiff (140 kPa) PDMS compared to soft (13 kPa) PDMS (**Extended Fig. 1**). While the percentage of contractile CMs was similar between soft and stiff substrates after two days in culture (61% vs. 57%) it was decreased by more than half on stiff substrates on day four (58% vs. 25%, **Fig. 2a**).

To analyze changes in the expression of genes associated to chromatin organization, we performed a comparative RNAseq analysis between cardiac cells plated on soft or stiff PDMS for four days. Of 114 annotated mouse histone genes^36^, we found 82 were expressed in our culture (**Extended Table 1**), of which 67 were more than 30% downregulated on stiff substrates (**Fig. 2b**). The expression of replication-dependent canonical histones was particularly reduced across all histone families which indicated that cardiac cell proliferation was inhibited on stiff substrates as reported previously^30^. However, we also observed downregulation of H1, H2a and H3 histone variants that replace canonical histones independent of replication and play a regulatory role in cell differentiation. H1^37^ and H2a^38^ variants have been implicated in the reprogramming of pluripotent stem cells and H3 variants have shown to play a role in neuronal development^39^ and cardiac hypertrophy^40^. This change of histone variant expression in stiff environments provided further evidence for a link between CM differentiation and chromatin organization.

Moreover, pathway analysis identified several signal transduction pathways involved in cell differentiation and cardiac signaling (**Extended Table 2**). Functional network grouping showed that pathways related to cell-ECM interactions were associated to MAPK (**Extended Fig. 2a**), an important pathway for cardiac development^41^ as well as epigenetic regulation^42^, which has been shown to influence chromatin positioning^43^. Gene ontology (GO) term enrichment analysis of differentially expressed (p<0.2) genes with the parent GO term *histone modification* (GO:0016570) revealed the child GO terms *histone methylation* (GO:0016571) and *histone acetylation* (GO:0016573) as significantly enriched (p=1.30E-9, p=5.50E-6). Closer investigation revealed that genes coding for the H3K9 methylases (aka methyltransferases) *Ehmt1*, *Setdb1* and *Eed* were among the most downregulated for cells cultured on stiff substrates (**Fig. 2c, Extended Table 3**). Methylation of H3K9 is associated with strong gene repression and chromatin condensation^44^ and has been shown to be crucial during cardiac development^45^ and maintenance^4^. RT-qPCR analysis validated the downregulation of H3K9 methylases, as well as downregulation of cardiac transcription factors and structural markers, while no change in gene expression was observed for the H3K9 demethylase *Kdm3a* (**Fig. 2d**) on stiff substrates compared to soft. *Eed* and *Ezh2* are also associated with the methylation of H3K27, in addition to H3K9 (**Extended Table 3**). Similar to H3K9, methylation of H3K27 is associated with gene repression and heterochromatin formation^44^; however, it has an opposing function during cardiac development and needs to be repressed for cardiac differentiation^46^. In accordance, PCR data showed that H3K27 demethylase *Kdm6a* was downregulated while H3K27-specific methylase *Ezh1* remained unchanged on stiff PDMS. Overall, gene expression analysis validated the inhibitory effect of stiff environments on CM differentiation and showed that the expression of histone variants and epigenetic enzymes, particularly those involved in H3K9 and H3K27 methylation, were altered in stiff environments.

### H3K9 and H3K27 Trimethylated Chromatin Shows Opposing Patterns of Reorganization Between Embryonic CMs and CFs During In Vitro Cultures

Similar to the nuclear organization in adult CMs, we observed an enrichment of chromatin at the nuclear border as well as a reduction in chromatin at the nuclear center when embryonic CMs were cultured for four days *in vitro* on soft (13 kPa) PDMS. We quantified overall chromatin enrichment by analyzing H2b-eGFP intensity over the relative distance to the nuclear center (0=center, 1=periphery) to verify this initial observation. A peripheral enrichment score was calculated as the ratio of the average intensity of the peripheral bin (0.85-0.95) divided by the center bin (0.05-0.15). Embryonic CMs, as identified via the formation of contractile myofibrils (**Fig. 3a, Extended Fig. 3a**), showed only minor accumulation of chromatin after two days in culture (**Fig. 3b-d**). However, after four days, chromatin was significantly enriched at the nuclear border (1.17 to 1.76). In contrast, we observed a decrease in peripheral chromatin enrichment from day two compared to day four in non-contractile CFs (1.09 to 0.98).

**Fig. 3.**
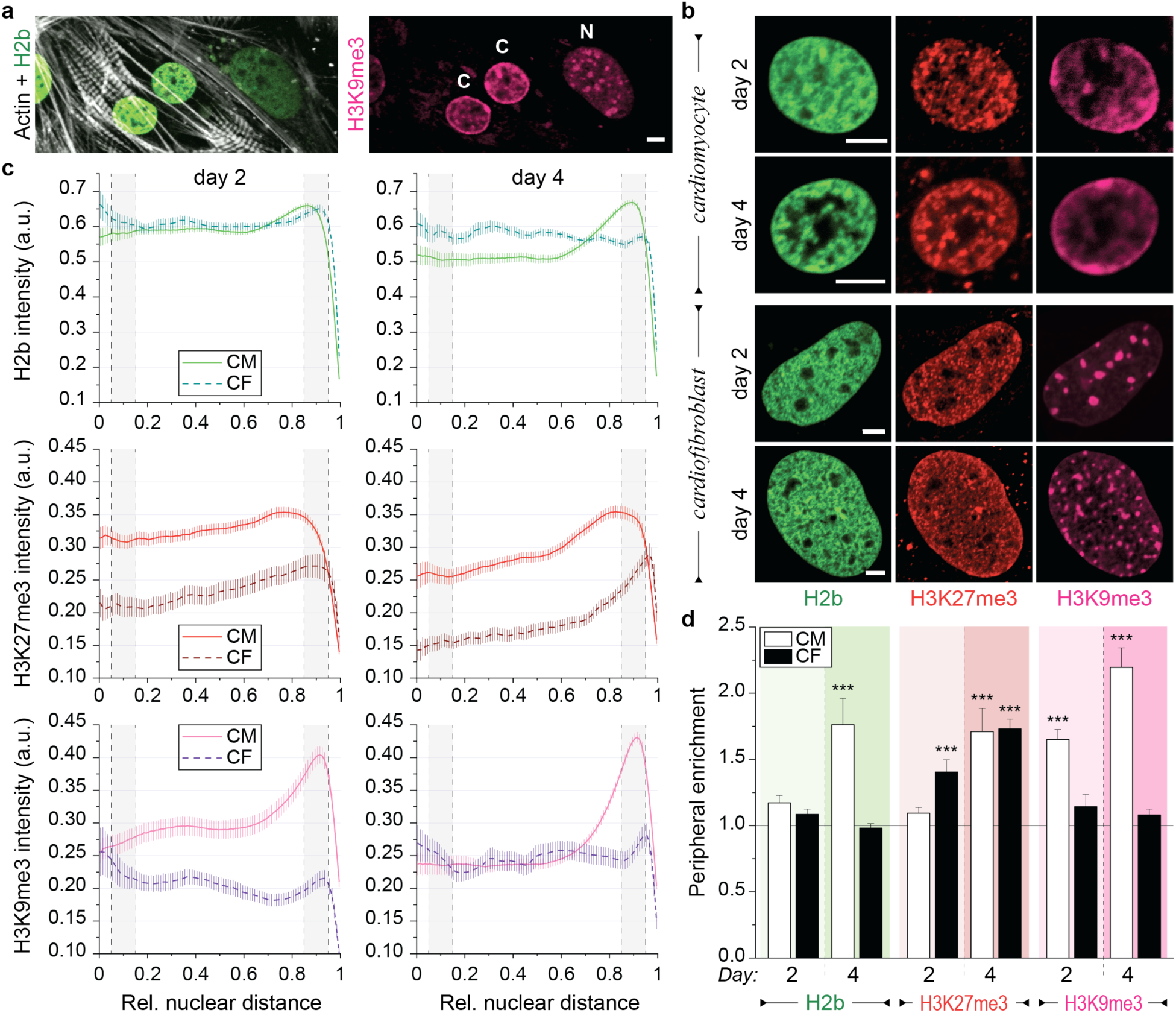
Contractile CMs and non-contractile CFs show opposing enrichment of H3K9 and H3K27 trimethylated chromatin *in vitro*. Embryonic cardiac cells were isolated from (E)18.5 H2b-eGFP embryo hearts and cultured on soft (13 kPa) PDMS substrates. **a)** After two days in culture, contractile CMs (C) with distinctly expressed myofibrils showed peripheral accumulation of H3K9me3-modified chromatin while non-contractile CFs (N) with actin fibers had a homogenous distribution of H3K9me3 clusters throughout the nucleus (see also Extended Fig. 3a). **b)** Cells were stained for H3K27me3 and H3K9me3 (and actin, see Extended Fig. 3b) and images of nuclei from CMs or CFs were acquired at day two or four of culture. **c)** Stained nuclei were analyzed for peripheral enrichment of overall chromatin (H2b) or epigenetically marked chromatin using a custom MATLAB code. Fluorescence intensity of each chromatin channel was analyzed with respect to its relative distance to the nuclear center (0=center, 1=periphery). Gray areas indicate the center bin (0.05-0.15) and the peripheral bin (0.85-0.95) used to calculate enrichment scores. SEM; n>60 from 5 exp. **d)** For each marker in each nucleus, an enrichment score was calculated as the quotient of intensity of the peripheral bin divided by the center bin. CMs, but not CFs, showed a shift of overall chromatin towards the nuclear border at day four. This was preceded by enrichment of H3K9me3-marked chromatin at both day two and four in CMs, while CFs showed enrichment of H3K27me3-modified chromatin instead; SEM; n>60 from 5exp.; T-test (HM=1): *** p<0.001. All scales=5 µm.

Gene expression analysis of cardiac cultures on stiff substrates associated to CM dedifferentiation revealed specific changes in the expression of enzymes involved in histone H3 methylation. Immunostaining of cardiac cultures showed that H3K9 trimethylated (H3K9me3) chromatin accumulated at the nuclear border in CMs while there was a homogenous distribution of H3K9me3 clusters throughout the nucleus of CFs (**Fig. 3a, Extended Fig. 3b**). Further quantification showed that H3K9me3-modified chromatin was significantly enriched at the nuclear border in CMs, but not in CFs on day two. This trend continued as we observed an increase in enrichment in CMs (1.65 to 2.20) compared to a slight decrease in CFs (1.14 to 1.08) (**Fig. 3d**). In turn, CFs showed increasing enrichment of H3K27me3-marked chromatin from days two to four (1.41 to 1.73). In CMs, peripheral enrichment of H3K27me3 occurred only on day four in conjunction with, and to the same extent as, overall chromatin. Furthermore, patterns of H3K27 methylation closely matched that of overall chromatin in CMs (**Fig. 3b**), suggesting that enrichment of H3K27me3-marked chromatin mainly resulted from overall chromatin rearrangement in CMs. In contrast, enrichment of H3K27me3 occurred in the absence of peripheral enrichment of overall chromatin in CFs. These results validated the contrary roles of H3K9 and H3K27 methylation in cardiac development, as embryonic CMs and CFs showed opposing patterns of enrichment over time. Furthermore, our results suggest that the trimethylation of H3K9 may play a role in guiding the observed chromatin reorganization during CM development, because the accumulation of H3K9me3-marked chromatin at the nuclear periphery preceded the accumulation of overall chromatin.

### Chromatin Reorganization is Abrogated in Stiffened Environments in Embryonic CMs In Vitro and Adult CMs In Vivo

H3K9me3-marked chromatin was enriched at the nuclear border of CMs whereas H3K27me3 was enriched at the border of CFs during our four-day *in vitro* culture model. To further investigate whether H3K9me3-associated chromatin reorganization is related to CM differentiation *in vitro*, we analyzed peripheral chromatin enrichment in embryonic CMs plated on soft (13 kPa) PDMS, stiff (140 kPa) PDMS, or tissue culture plastic (TCP, >1 GPa). Surprisingly, CMs plated on TCP showed a higher overall chromatin enrichment at the nuclear periphery compared to cells on either PDMS substrates (**Fig. 4a-c**) on day two. However, on day four peripheral enrichment on stiff substrates declined and CMs on soft PDMS showed a higher peripheral accumulation of overall chromatin compared to both stiff PDMS and TCP (1.76 vs. 1.12 and 1.12). In accordance, H3K9me3-marked chromatin was equally enriched for CMs on any substrate on day two, whereas on day four enrichment was higher on soft PDMS compared to both stiffer substrates and was higher for stiff PDMS compared to TCP (2.20 vs. 1.87 vs. 1.25). While there was no difference in enrichment on day two, CMs on soft PDMS had higher H3K9me3 intensities at any distance compared to cells on stiffer substrates (**Fig. 4b**) in accordance with decreased expression of H3K9 methyltransferases observed on stiff PDMS (**Fig. 2c**). H3K27me3-marked chromatin was slightly more enriched for CMs on stiff PDMS on day two compared to cells on soft PDMS. After four days, overall enrichment of H3K27me3-modified chromatin increased for all conditions and was highest in cells plated on the soft substrates; however, H3K27me3 enrichment was higher compared to overall chromatin on stiff substrates similar to observations in non-contractile CFs. Substrate stiffness moderately affected chromatin organization in CFs with peripheral enrichment being low for overall and H3K9me3-marked chromatin and high for H3K27me3-modified chromatin throughout the four-day culture period (**Extended Fig. 4a-c**), suggesting that CMs are more sensitive to substrate stiffness with regard to chromatin reorganization.

**Fig. 4.**
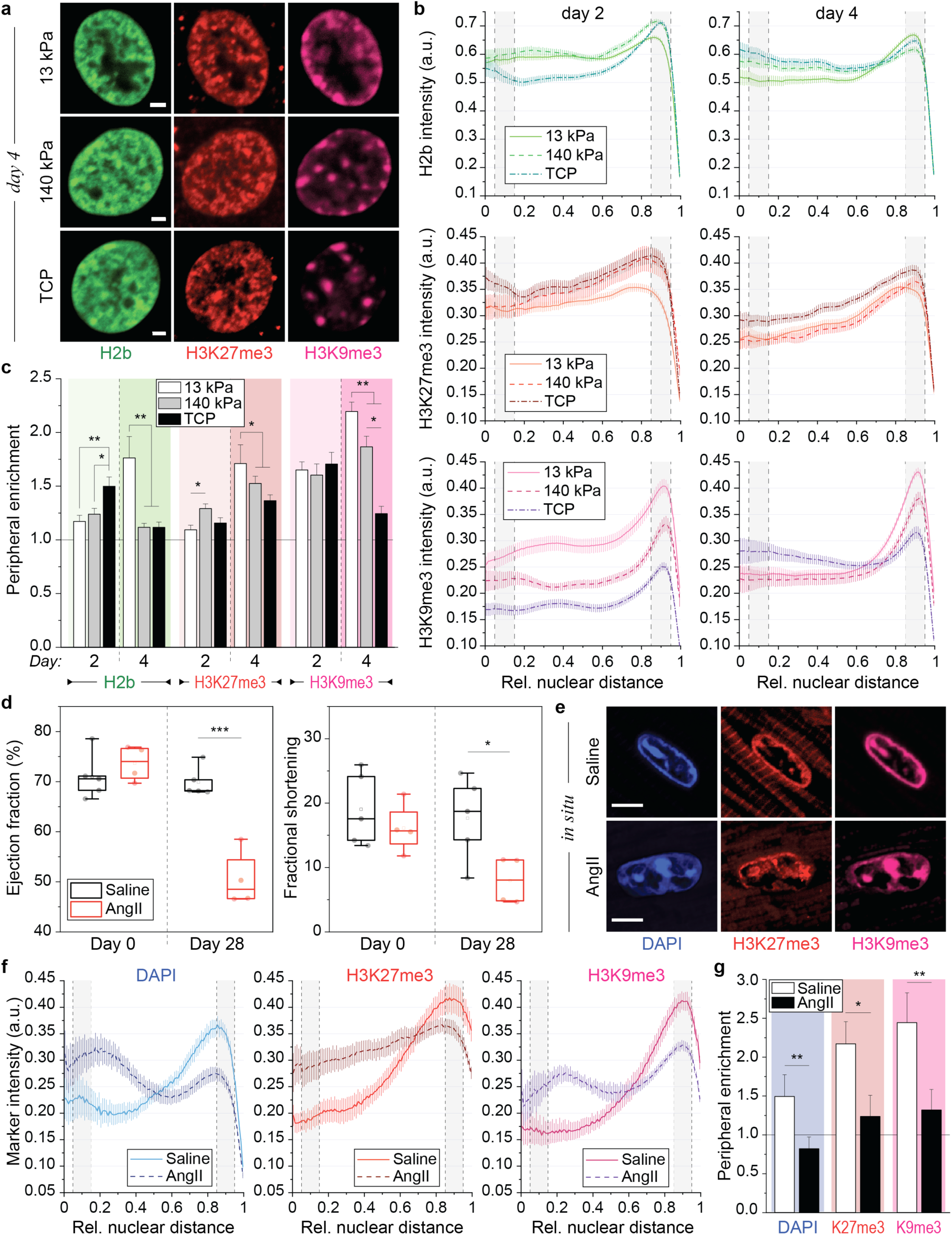
Chromatin reorganization is inhibited in embryonic CMs cultured on stiff substrates *in vitro* and abrogated in adult CMs during hypertrophy *in vivo*. **a)** Embryonic cardiac cells were isolated from (E)18.5 H2b-eGFP embryo hearts and cultured on either soft (13 kPa) PDMS, stiff (140 kPa) PDMS or TCP for two or four days after which nuclei were stained for H3K27me3 and H3K9me3 as well as actin to distinguish CMs from CFs. **b)** CM nuclei were evaluated for peripheral enrichment of overall chromatin (H2b) or epigenetically marked chromatin using a custom MATLAB code that analyzed marker intensity with respect to its relative distance to the nuclear center (0=center, 1=periphery). Gray areas indicate center and peripheral bin; SEM; n≥60 from 5 exp. **c)** Enrichment scores for each chromatin marker were calculated as the quotient of intensity of the peripheral bin (0.85-0.95) divided by the center bin (0.05-0.15). Enrichment of overall and H3K9me3-marked chromatin was abrogated on day four in nuclei of cells plated on stiff PDMS and TCP compared to soft PDMS. Note: 13 kPa data same as CM data in Fig. 3d; SEM; n≥60 from 5 exp.; 1W-ANOVA: * p<0.05, ** p<0.01, *** p<0.001. **d)** Mice treated with angiotensin II (AngII, n=4), to induce cardiac hypertrophy, showed reduced ejection fraction and fractional shortening after 28 days of treatment compared to day 0, while saline receiving control mice (n=5) showed no difference; T-test: * p<0.05, *** p<0.001. **e)** After 28 days, hearts were harvested and stained for H3K27me3 and H3K9me3. DAPI was used as DNA counterstain. **f, g)** Immunostained cardiac sections of hypertrophic (AngII) or control mice (Saline) were analyzed for peripheral enrichment of overall chromatin (DAPI) or epigenetically marked chromatin using a custom MATLAB code and enrichment scores were calculated. Enrichment of overall and methylated chromatin was abrogated in cardiac nuclei of hypertrophic mice while control mice showed a mature cardiac phenotype; SEM; n≥40 from 4 (AngII) or 5 (saline) exp.; T-test: * p<0.05, ** p<0.01. All scales=5 µm.

Hypertrophy leads to an increase in cardiac stiffness and CM dedifferentiation^4,5^. To validate our findings *in vivo*, we analyzed CM nuclei in mice that received angiotensin II (AngII) for 28 days to induce hypertrophy. Control mice received saline over the same period. Reduction in ejection fraction and fractional shortening during the treatment period confirmed cardiac performance decline in mice receiving AngII, but not saline (**Fig. 4d**). CM nuclei of hypertrophic mice showed reduced enrichment of overall (DAPI, 0.82 vs. 1.49), H3K27me3 (1.24 vs. 2.17) and H3K9me3-marked chromatin (1.32 vs. 2.44, **Fig. 4e-g, Extended Fig. 4d**) similar to CM nuclei cultured on stiff substrates *in vitro*. Together these results further supported a link between H3K9 trimethylation and chromatin reorganization during CM differentiation and suggested that mechanical environments play an important role to establish and retain the nuclear phenotype of adult CMs.

### LINC Complex Disruption Inhibits Reorganization of H3K9 Methylated Chromatin in Embryonic CMs, but Does Not Regulate Histone (De)Methylase Expression

We observed enrichment of H3K9me3-marked chromatin in contractile CMs, but not in non-contractile CFs. This enrichment was inhibited in stiffened environments that lead to a reduction in CM contractility^28^. In accordance, analysis of nuclear deformation recorded during CM contraction showed that bulk linear strain and translational movement of nuclei were reduced in cells plated on stiff PDMS and TCP over the four-day culture period (**Extended Fig. 5a**). CM nuclei are connected to the Z-disks of myofibrils via LINC complexes^13^, which are crucial for nuclear mechanotransduction^14^ and cardiac development^13,16,17^. To further investigate the link between nuclear deformation and chromatin reorganization, we disrupted LINC complexes in embryonic CMs by overexpressing a truncated nesprin-3 protein that contained the transmembrane and KASH (Klarsicht, ANC-1, Syne Homology) domain (Δsyne-K3) but lacked cytoskeleton binding domains (**Fig. 5a**). Because KASH domains are highly conserved between species and nesprins, this construct competes with all nesprin isoforms for SUN connections^47–49^. The control vector expressed a protein that was identical to Δsyne-K3 but lacked the KASH domain needed for LINC complex integration (Δsyne-CTL). Both vectors also expressed fluorescently tagged α-actinin 2 to identify infected cells.

**Fig. 5.**
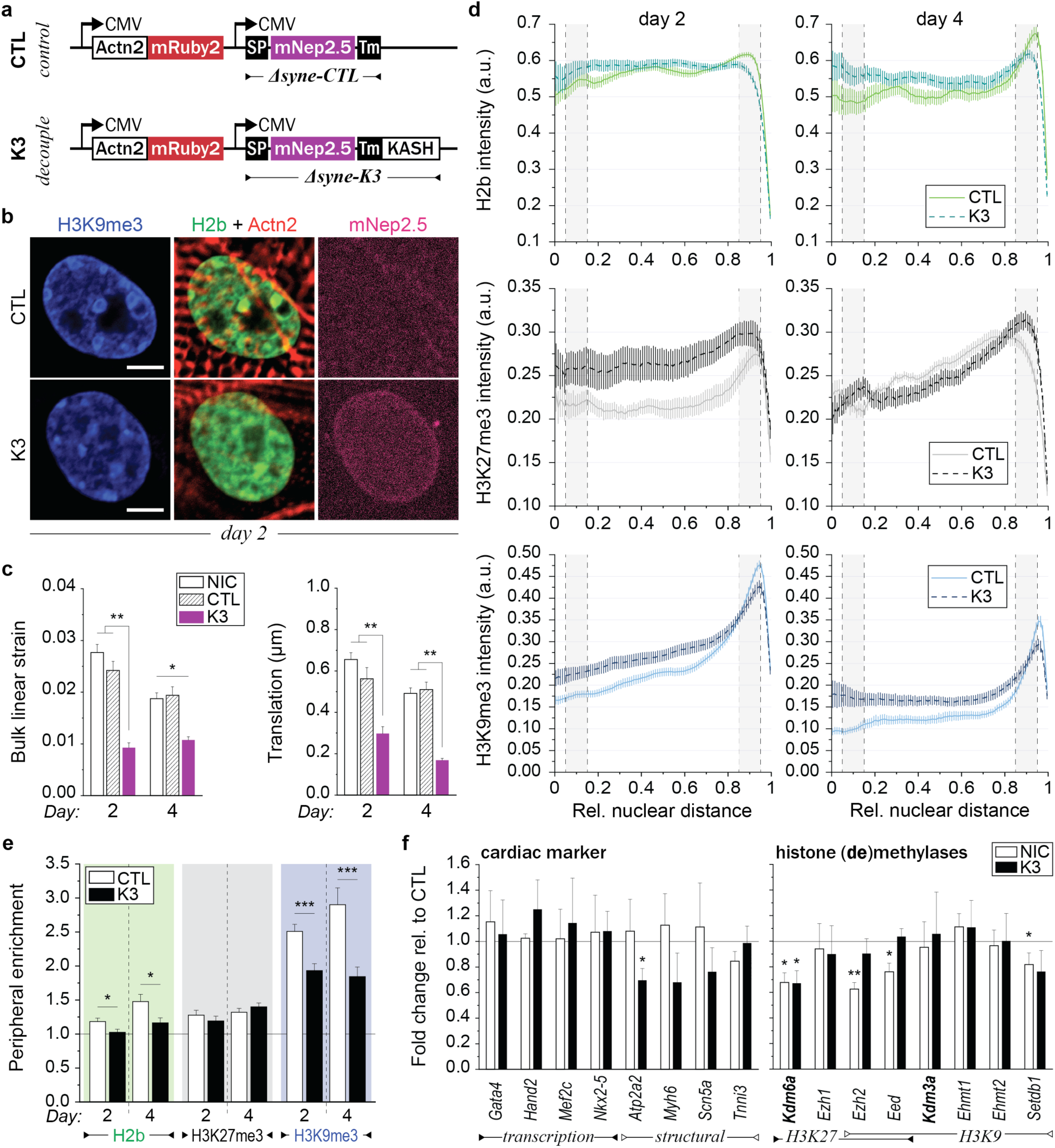
LINC complex disruption abrogates rearrangement of H3K9 methylated chromatin in CMs but does not affect epigenetic enzyme expression. Embryonic cardiac cells were isolated from (E)18.5 H2b-eGFP embryo hearts and cultured on soft (13 kPa) PDMS substrates. **a)** Illustration of adenoviral vectors for LINC complex disruption. The decoupling vector (K3) expressed a truncated nesprin-3 composed of the transmembrane (TM) and the KASH domain tagged with mNeptune2.5 (mNep2.5) to track nuclear membrane integration. The control vector (CTL) lacked the KASH domain necessary for binding to SUN within LINC complexes. Fluorescently tagged α-actinin 2 (*Actn2*) was expressed to identify infected CMs. **b)** Cells were infected with either vector on day one and stained for H3K9me3 (or H3K27me3, see Extended Fig. 5c) on day two (shown) or day four. The truncated nesprin construct integrated successfully into the outer nuclear membrane of infected CMs (mNep2.5) while no distinct localization was observed for the control vector. Myofibril formation was disrupted in decoupled CMs, particularly around the nucleus, which indicated successful decoupling of nuclei from the cytoskeleton (see also Extended Fig. 5d); scale=5 µm. **c)** Image series of nuclei during CM contractions were recorded on day two (24h post infection) and day four and bulk linear strain and translational movement of nuclei were determined. Decoupled nuclei (K3, n=32) showed lower bulk linear strain and translational movement compared to cells infected with the control vector (CTL, n=32) or non-infected control cells (NIC, n=67); SEM; from 4 exp.; 1W-ANOVA: * p<0.05, ** p>0.01. **d, e)** Infected cells were stained for H3K9me3 or H3K27me3 on day two or four and peripheral enrichment of chromatin was analyzed using a custom MATLAB code. Decoupled cells (K3) showed abolished enrichment of overall and H3K9me3-marked chromatin compared to infected control cells (CTL) while H3K27me3-marked chromatin was similarly enriched; SEM; n=35 from 3 exp.; T-test: * p<0.05, *** p<0.001. **f)** Gene expression analysis of decoupled (K3) or non-infected control cells (NIC) compared to infected control cells (CTL). Expression of structural, but not transcriptional, cardiac genes was reduced in decoupled cells, while expression of histone methylating and demethylating genes remained largely unchanged except for a decrease of *Kdm6a*; SD; n=4; T-test: * p<0.05, ** p<0.01.

Cardiac cultures were plated on soft (13 kPa) PDMS and infected on day one of culture. CMs that successfully integrated Δsyne-K3 at the nuclear border 24 hours after infection (day 2) had disorganized sarcomere fibers, particularly around the nucleus (**Fig. 5b, Extended Fig. 5b**, **Extended Videos 1-3**), diminished localization of nesprin-1 at the nuclear periphery (**Extended Fig. 5b**), and decreased nuclear bulk linear strain and translational movement during contraction (**Fig. 5c**), all of which indicated successful LINC complex disruption. Enrichment towards the nuclear border was significantly impaired for overall (H2b) and H3K9me3-modified chromatin in decoupled CMs compared to control cells, while H3K27me3 showed no distinct difference in enrichment (**Fig. 5b, d and e and Extended Fig. 5c**). Notably, overall H3K9me3 intensities were similar between decoupled and control cells, while H3K27me3 intensities were slightly elevated after decoupling (**Fig. 5e**) similar to results on stiff substrates. RT-qPCR analysis revealed that gene expression of H3K9 and H3K27 histone (de)methylases was largely unchanged in decoupled cells, except for the downregulation of H3K27 demethylase *Kdm6a* (**Fig. 5f**), suggesting that LINC-associated nuclear mechanosensitive pathways do not regulate the expression of epigenetic mediators. Expression in non-infected control cells (NIC) was altered compared to infected control cells, indicating that adenovirus transfection affected the expression of epigenetic enzymes as previously reported^50^. In addition to epigenetic modifiers, decoupling also did not affect the expression of cardiac-specific transcription factors while structural cardiac markers were partially downregulated. Together these results showed that the disruption of LINC complex connections from the cytoskeleton to the nucleus inhibited the reorganization of H3K9me3-modified chromatin in CMs but did not affect the expression of cell-fate mediators such as transcription factors or epigenetic enzymes.

### H3K9 Trimethylated Chromatin Is Co-Localized with Myofibrils in Embryonic CMs

We observed that the reorganization of H3K9me3-modified chromatin towards the nuclear periphery was inhibited for CMs in stiff environments and after LINC complex disruption, both of which reduced nuclear deformation, which suggested a potential mechanosensitive feedback between myofibrils and the nucleus. Analyzing the shortening of myofibrils of CTL infected CMs via α-actinin 2-mRuby2 expression further validated that CM contractility was abrogated on stiff (140 kPa) compared to soft (13 kPa) PDMS as overall sarcomere (S) and A-band (A) shortening were reduced by more than half (−0.171 vs. −0.077 µm, −0.185 vs. −0.060 µm), and Z-disks (Z) went from extending to shortening (0.015 vs. −0.032 µm; **Fig. 6a-c**). In accordance with abrogated Z-disk extension, stretch-induced tyrosine-410 phosphorylation of p130Cas^51^, a mechanosensitive focal adhesion protein^52^ found within Z-disk lattice in CMs^53^, was reduced on stiff PDMS as well (**Extended Fig. 2b**). Contractility was also abrogated in decoupled CMs infected with K3, albeit to a lesser extent than CMs on stiff PDMS (**Fig. 6c**), which highlighted the role of LINC complexes in myofibril formation as recently reported^54^.

**Fig. 6.**
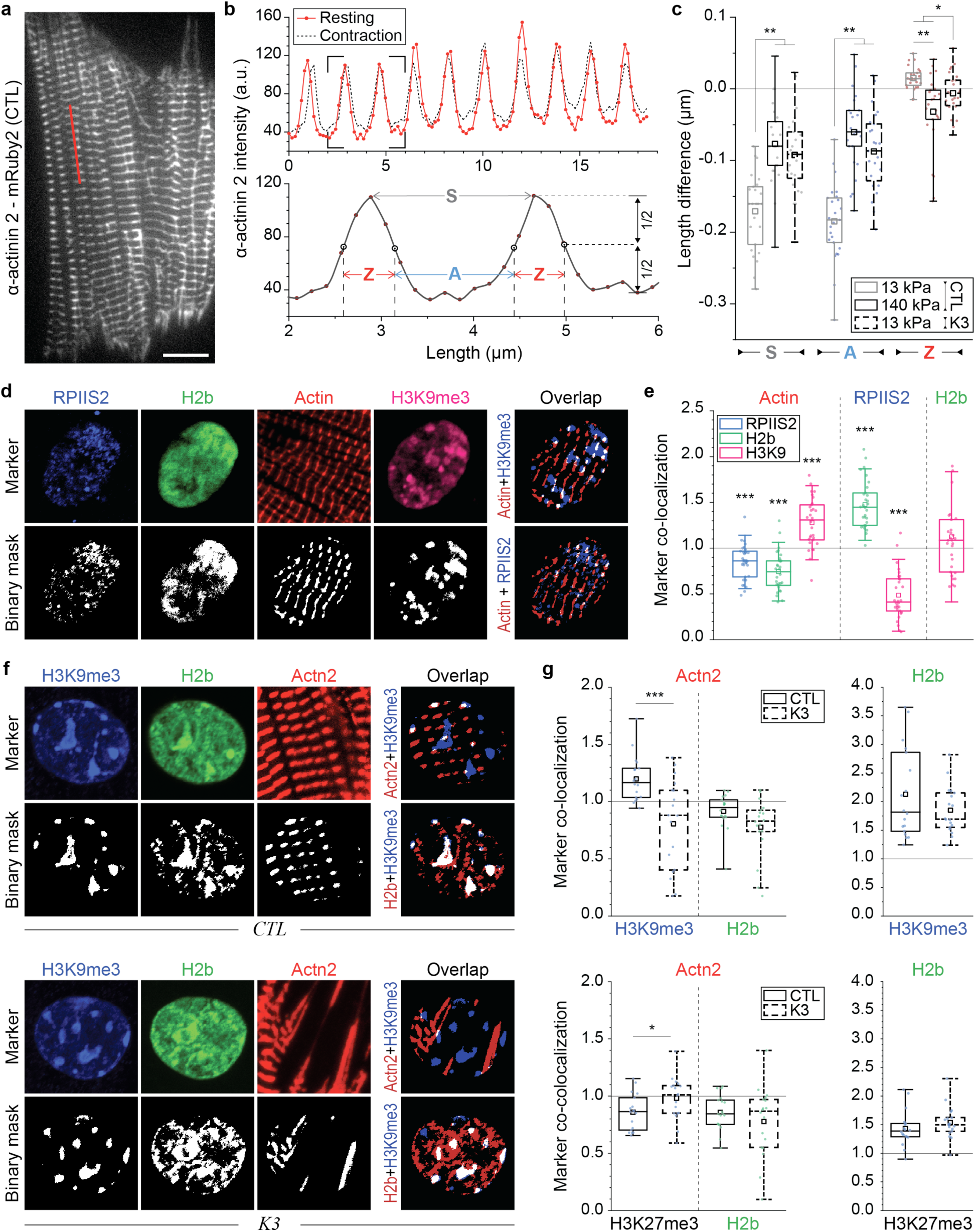
Co-localization of H3K9me3-marked chromatin with myofibrils is abrogated in CMs after LINC complex disruption. Embryonic cardiac cells were isolated from (E)18.5 H2b-eGFP embryo hearts. **a)** CMs were infected with the adenoviral decoupling vector K3 or control vector CTL (shown) on day one and image series of myofibril contraction were recorded on day four via fluorescently tagged α-actinin 2; scale=10 µm. **b)** Top: α-actinin 2 intensity profile, as indicated by a red line in a), before (resting) and during contraction. Bottom: Close-up of two intensity peaks. Analysis of intensity profiles was used to determine the difference in length of overall sarcomeres (S), A-bands (A) and Z-disks (Z) during contraction. **c)** Control infected CMs on stiff PDMS (140 kPa, CTL) and decoupled CMs on soft PDMS (13 kPa, K3) showed inhibited contraction as overall sarcomere and A-band shortening as well as Z-disk extension was abrogated compared to control infected CMs on soft PDMS (13 kPa, CTL); n=25 from 5 exp.; 1W-ANOVA: * p<0.05, ** p<0.01. **d, e)** After four days in culture on soft PDMS, CMs were stained for regions of active transcription (RPIIS2), sarcomeric I-bands (actin) and H3K9me3. Marker channels were binarized and a co-localization score was calculated for each marker pair as the percentage of overlapping pixels divided by their independent probability to overlap (1=chance). H3K9me3-marked chromatin showed above chance associating with actin while regions of active transcription and overall chromatin did not; n=30 from 3 exp.; T-test (HM=1): *** p<0.001. **f, g)** CMs on soft PDMS were infected with K3 or CTL on day one and stained for H3K9me3 (shown) or H3K27me3 (Extended Fig. 6b) on day four to analyze marker overlap. Co-localization of H3K9me3 with α-actinin 2 containing Z-disks (Actn2) was abrogated after LINC complex disruption while co-localization of H3K27me3 was increased; n=18 from 3 exp.; T-test: * p<0.05, *** p<0.001.

To further inquire whether there is a direct association of H3K9me3-marked chromatin with myofibrils, we quantified the relative overlap of different chromatin markers with myofibrils in CMs cultured for four days on soft (13 kPa) PDMS. In addition to H3K9me3 and actin, cells were stained for serine-2 phosphorylated RNA-polymerase II (RPIIS2) as a control (**Fig. 6d**) since actively transcribed chromatin^55^ is expected to be exclusive with suppressed H3K9me3-modified chromatin. Z-stacks of stained CMs were acquired and basal Z-slices, where myofibrils were primarily located in our *in vitro* cultures (**Extended Fig. 4a**), were analyzed for marker overlap. A marker co-localization score was calculated by determining the percentage of overlapping pixels between two binarized marker channels normalized over their independent probability to overlap, with a score of 1 representing marker co-localization by chance (see Methods for details). H3K9me3-marked chromatin showed a higher than chance association with actin containing I-bands (1.28) while overall (H2b) and actively transcribed chromatin had lower than chance co-localization scores (0.74 and 0.85, **Fig. 6e**). The low association with overall chromatin is likely an effect of actin pushing chromatin out of the *z*-plane (**Extended Fig. 6a**). As expected, transcribed chromatin areas had a high coincidence score for overall chromatin but a low score for H3K9me3 (1.48 vs. 0.49) while the association of overall chromatin with H3K9me3 did not significantly deviate from chance (1.12, p=0.165).

To further analyze marker co-localization after LINC complex disruption, we infected cardiac cells with the decoupling vector K3, or control vector CTL, on day one and stained for H3K9me3 or H3K27me3 on day four (**Fig. 6f-i**, **Extended Fig. 6b**). We observed an above chance coincidence score for H3K9me3-marked chromatin with the Z-disk protein α-actinin 2 (Actn2), which was significantly decreased below chance after decoupling (1.20 vs. 0.81). In contrast, coincidence scores of H3K27me3 with Actn2 were slightly increased after decoupling (0.86 vs. 0.98), again highlighting the inverse relationship between H3K9me3 and H3K27me3 modifications in CM development. No significant difference in overlap of overall chromatin and Actn2 was observed in either decoupled or control cells; however, both infected groups showed an increased association of H3K9me3 with overall chromatin compared to uninfected cells, again suggesting a potential influence of adenoviral infections on epigenetic regulation. Overall, these findings provided evidence that H3K9me3-marked chromatin is associated with myofibrils in CMs, but only when connected to the nucleus via LINC complexes, further consolidating a link between nuclear mechanosensation and chromatin reorganization.

### H3K9 Trimethylated Chromatin Domains are Localized to Intranuclear Subregions with Elevated Tensile Strains During CM Contractions

Our results suggested a link between myofibril-mediated nuclear deformation and peripheral enrichment of H3K9 trimethylated chromatin as well as an association of H3K9me3 with myofibrils. Analysis of CM nuclei during contractions further confirmed this association as dense, H3K9me3-rich heterochromatin clusters had higher translational movement compared to the overall nucleus (**Extended Fig. 6c**), To investigate the link between nuclear strain and chromatin reorganization in CMs, we performed an in-depth analysis of the strain occupancy for different chromatin types during CM contraction. For that we utilized a recently published method, termed *deformation microscopy*^31,32^, which generates high-resolution spatial strain maps from an undeformed template and a deformed target image (**Fig. 7a**). Image series of nuclear deformations were recorded in contracting CMs on day two after which cells were fixed, stained, and co-registered to chromatin markers. CMs recorded for live imaging were relocated, imaged and the common H2b-GFP channel was used to register intranuclear strain maps with chromatin marker maps to calculate strain values for individual chromatin markers or no marker (unassigned, **Fig. 7b**). Strain values were normalized to the average strain of each nucleus to enable comparison between cells.

**Fig. 7.**
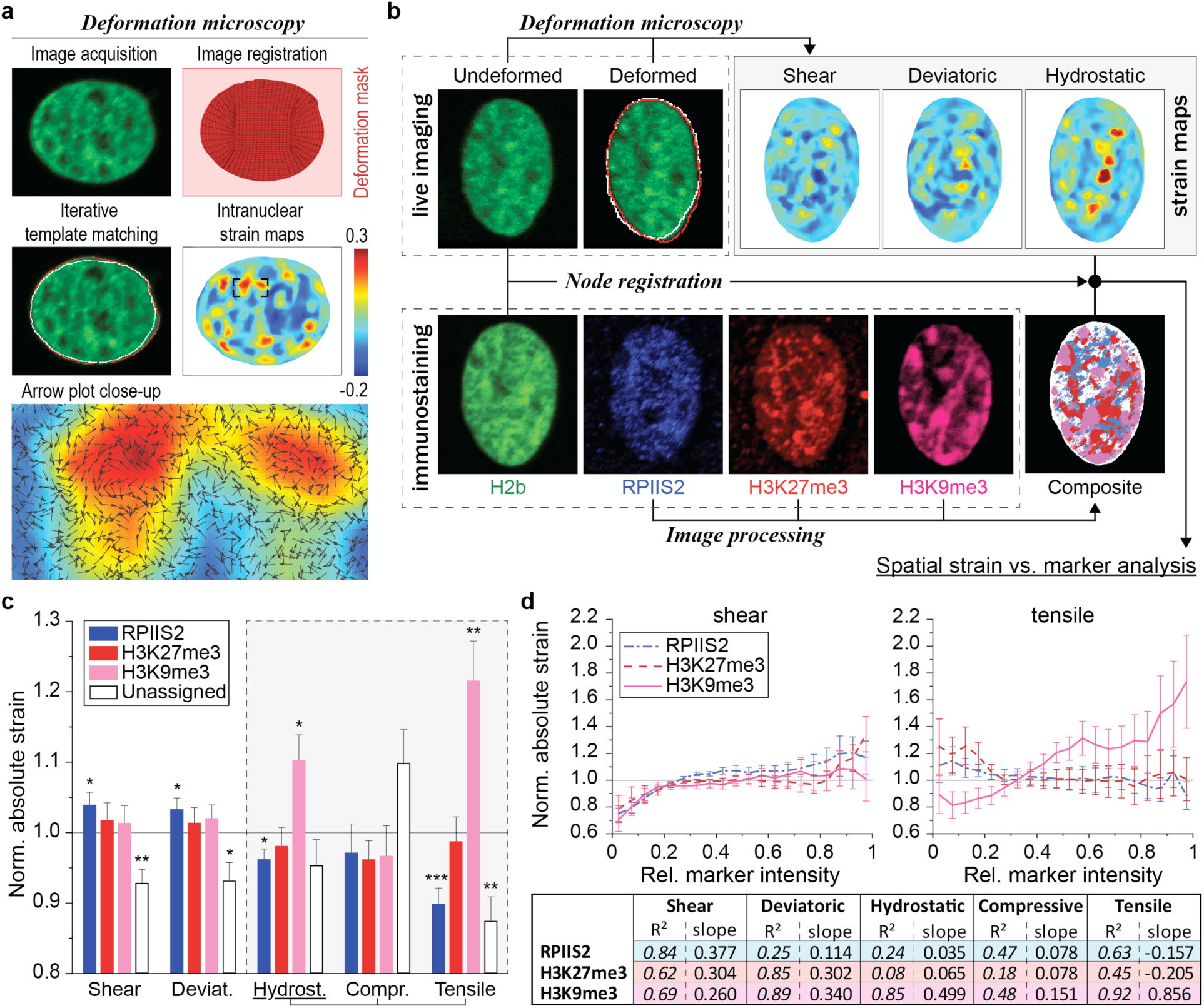
H3K9me3-marked chromatin is localized to intranuclear regions with elevated tensile strains during CM contractions. **a)** Illustration of *deformation microscopy* to generate high-resolution strain maps from image data. Image series of H2b-eGFP CM nuclei were acquired during contraction. Frames of undeformed nuclei during diastolic resting (white outline) were iteratively registered and warped to match nuclear image frames during peak contraction (red outline). Arrow plot shows a close-up of the resulting intranuclear strain map for hydrostatic strains. **b)** Flowchart of spatial strain vs. marker analysis using *deformation microscopy*. Image series of CM nuclei were recorded on day two to calculate intranuclear strain maps. After, CMs were stained for H3K9me3, H3K27me3 or actively transcribed chromatin (RPIIS2), relocated and imaged again. Spatial nodes between both datasets were registered via the common H2b channel and strain occupancy for each marker was analyzed. **c)** Composite analysis of nuclear strains over chromatin assigned to one marker or none of the markers (unassigned, white areas in composite image in b). Strains were normalized to the average of each nucleus. H3K9me3-marked chromatin showed above average association with tensile hydrostatic strain; SEM; n=20 from 5 exp.; T-test (HM=1): * p<0.05, ** p<0.01, *** p<0.001. **d)** Continuous analysis of intranuclear strains over chromatin marker intensities. Top: Strain vs. intensity plot for shear and tensile hydrostatic strain (see Extended Fig. 7 for other strains). Bottom: Linear regression summary of strain vs. intensity plots. Highest R^2^ and steepest slope were observed for tensile hydrostatic strains over H3K9me3 intensities; SEM.

H3K9me3-marked chromatin areas experienced above average absolute hydrostatic (changes in volume) strain magnitudes (1.103), but not shear or deviatoric strains (1.014 and 1.021, **Fig. 7c**). Interestingly, subdividing hydrostatic strain values into tensile (positive) and compressive (negative) revealed that this trend arose from tensile strains alone (1.216) as no significant difference was observed for compressive hydrostatic strains (0.967) between any of the investigated chromatin features. In contrast, transcribed chromatin regions (RPIIS2) showed lower absolute hydrostatic strain (0.962), a trend that was again augmented for tensile hydrostatic strain (0.899), while no significant difference was observed for H3K27me3 for any of the strains investigated (p>0.177). Linear regression analysis of chromatin marker intensities over intranuclear strains further showed the highest degree of correlation (R^2^=0.923) and the highest slope for tensile strains over H3K9me3-intensities (**Fig. 7d** and **Extended Fig. 7a**). Analysis of intranuclear strains over the distance to the nucleus center showed that strains generally declined towards the periphery, excluding the possibility that H3K9me3-marked chromatin and hydrostatic strains simply coincide at the nuclear border (**Extended Fig. 7b**).

Moreover, analysis of marker intensities over their angle with respect to the nuclear center revealed that H3K9me3 localization peaked within ±30° in the direction of nuclear translation; however, only in nuclei with a tensile loading mode (**Extended Fig. 8**). In contrast, nuclei with a shortened major axis during contraction showed diminished H3K9me3 occupancy at the angle of nuclear translation while H3K27me3 and RPIIS2 showed no observable trend for any loading mode. Together, these findings provided further support for a link between the reorganization of epigenetically modified chromatin and contraction-mediated nuclear deformation in CMs and connects previous observations of the interactions between environment (substrate stiffness), cell differentiation and cell-type specific nuclear architecture into a new model of nuclear mechanosensation (**Fig. 8**).

**Fig. 8.**
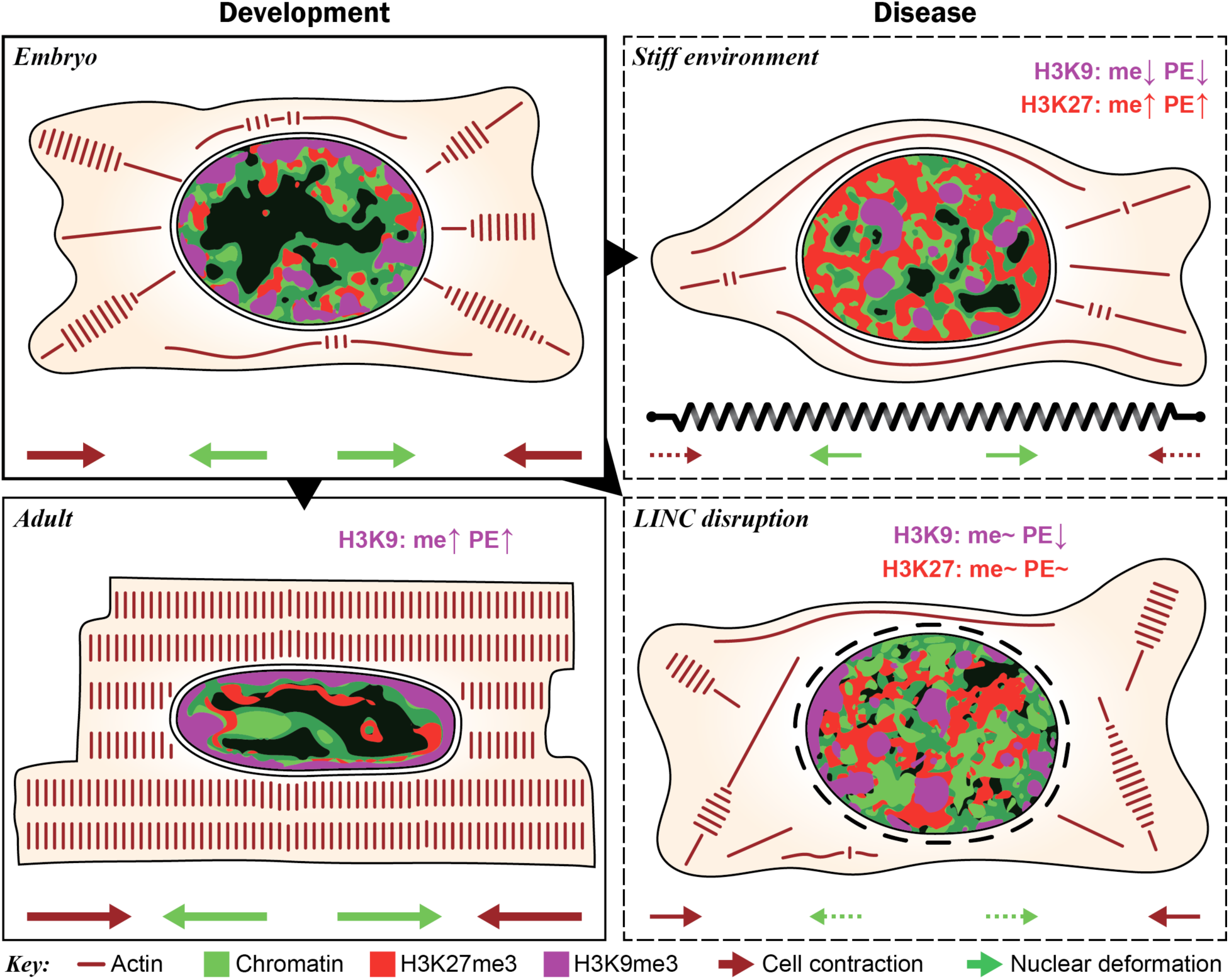
Summary of chromatin reorganization and epigenetic regulation during cardiac development and disease. During development, myofibril-mediated cell contraction and subsequent nuclear deformation of embryonic CMs leads to increased H3K9 trimethylation (me) and peripheral enrichment (PE) of H3K9me3-modified chromatin. This presumably stabilizes cardiac gene expression by anchoring suppressed non-cardiac genes to the periphery to prevent reactivation. Stiffening of the cardiac micro-environments inhibits cell contraction and nuclear deformation resulting in increased trimethylation and peripheral enrichment of H3K27 instead of H3K9. Abrogation of nuclear strain transfer through disruption of LINC complexes inhibited chromatin reorganization while regulation of methylation remained largely unchanged, suggesting that nuclear mechanosensation primarily affects chromatin reorganization.

## DISCUSSION

We showed that CMs establish a cell-type specific nuclear architecture during development that is characterized by a relocation of chromatin from the interior to the nuclear periphery. Our findings suggest that tensile strains transferred from myofibrils to the nucleus via LINC complex connections guide the rearrangement of epigenetically suppressed chromatin to the nuclear periphery. Stiff environments, which inhibited contraction and decreased nuclear deformation, or disruption of LINC complexes, profoundly disturbed the establishment of this architecture in embryonic stages or its maintenance in adults. We additionally observed opposing trends for the relocation of H3K9me3 and H3K27me3-modified chromatin as peripheral enrichment of H3K9me3 proceeded that of overall chromatin in embryonic CMs while early relocation of H3K27me3 was indicative for the formation of a non-contractile fibroblast cell-type. Taken together, our results provide further support for a link between nuclear architecture and cell differentiation and suggest that environmental cues can influence cell fate by affecting epigenetic regulation and reorganization of epigenetically modified chromatin (**Fig. 8**). This work therefore connects previously observed nexuses between substrate properties, cell fate, and nuclear mechanics into one conceptual framework that highlights the role for nuclear mechanosensitive pathways in guiding and stabilizing cell fate determination through spatial chromatin organization. Our data from LINC complex disruption experiments indicated that epigenetic regulation is mediated independently of nuclear strains, hence the interplay between cells and their environment is likely to be a more complex interaction between nuclear and cellular mechanosensors.

### Potential Roles of Chromatin Organization in CMs

The observed reorganization of H3K9me3-marked chromatin during CM differentiation might function to stabilize differential gene expression by segregating repressed non-cardiac genes to areas of low activity, like the nuclear envelope^56^, to prevent accidental reactivation. In this study, RNAseq and PCR gene expression analysis revealed that H3K9 methylases *Ehmt1*, *Ehmt2*, *Setdb1,* and *Eed* were particularly downregulated on stiff substrates together with cardiac differentiation markers. Recent studies have also shown that the expression of *Ehmt1/2* was decreased in the heart of hypertrophic rats and that the reactivation of cardiac fetal genes such as *Myh7*, *Nppa*, and *Nppb* was associated with the loss of H3K9 methylations at those genes^4^. Furthermore, inhibition of *Ehmt1/2* was sufficient to induce hypertrophy while stabilization of *Ehmt1/2* expression counteracted effects of hypertrophy. Similarly, targeted deletion of H3K9 demethylase *Kdm4a* in the heart led to attenuated hypotrophy in mice^5^. Here we observed that H3K9me3-marked chromatin relocated back towards the nuclear center in mature CMs of hypertrophic mice, supporting the hypothesis that peripheral segregation of chromatin stabilizes its suppression. Interestingly, mechanical decoupling of CM nuclei through LINC complex disruption only minorly affected the expression of H3K9 (de)methylating enzymes or cardiac transcription factors. It is likely that CMs integrate mechanical signals from different pathways to achieve reliable and robust cell differentiation. One such candidate, identified here via RNAseq and validated through western blot analysis, was p130Cas. This mechanosensitive focal adhesion protein^52^ is found within the sarcomeric Z-disk lattice in CMs^53^, hence making it a suitable candidate to sense substrate stiffness and influence epigenetic regulation via Rap1 and MAPK signaling (**Extended Fig. 2**). One study also showed that p130Cas knockout mice die *in utero* due to abnormal heart growth, further linking p130Cas to cardiac development^57^.

In addition to stabilizing cardiac gene transcription, the CM nuclear architecture might also have a structural function as has been reported for other types of cells^24,25,58^. The particularly strong accumulation of heterochromatin at the nuclear border might play a protective role as heterochromatin has been shown to increase nuclear rigidity independent of lamins^59^. Chromatin void cavities could provide a protective environment for gene transcription in cells that must endure high and constant cyclic mechanical stresses. This is supported by our observation that nuclear areas of actively transcribed chromatin experienced lower (tensile) hydrostatic strains (**Fig. 7**).

### Potential Role of Nuclear Mechanosensation in CMs

The role of nuclear mechanotransduction has long been debated. While it has been shown that LINC complexes are involved in mechanosensitive gene regulation^60^, no distinct mechanism has been reported so far. We provided evidence that nuclear strains transferred from myofibrils via LINC complexes guide the reorganization of H3K9me3-modified chromatin in CMs. In contrast, the formation of a non-contractile CFs was associated with the reorganization of H3K27me3-marked chromatin instead. Our data suggests that CMs may use nuclear deformation as a feedback for differentiation by consolidating the suppression of H3K9me3-marked non-cardiac genes through peripheral segregation as described above. This mechanism would be beneficial during development as it ensures that only CMs with high contractility mature, while others become fibroblasts or undergo apoptosis. Neurons are known to use similar activity-based mechanisms during development in which their initial state of differentiation is reinforced through functional fidelity^61^. Further investigation will show whether this nuclear-feedback is used by other load-bearing cells as well.

### Potential Mechanisms of Mechanosensitive Chromatin Reorganization

How nuclear strains could guide chromatin rearrangement is unclear, but it is likely to involve nuclear motor proteins. For example, nuclear myosin 1 (NM1) has been shown to be necessary for chromatin relocation in response to serum activation^62^ and DNA damage^63^. After transport, nuclear envelope transmembrane proteins (NET) could anchor chromatin to the nuclear envelope. NETs have been shown to be tissue specific and might also recognize epigenetic modification of chromatin^20,64,65^. Nuclear motor proteins and NETS are poorly characterized, and more research is needed to understand their role in the nucleus.

The strict association of H3K9me3 with tensile, but not compressive, hydrostatic strains indicated the involvement of stretch-activation. It has been argued that LINC complexes are the nuclear analog to stretch-sensitive focal-adhesions at the cell membrane^66^, and research on isolated nuclei has shown that LINC complexes are necessary for stretch-induced nuclear mechanotransduction^14^. Similarly, we found in this study that LINC complexes play a role for chromatin reorganization in CMs. Direct evidence for stretch-sensitivity of LINC complex components (nesprins and SUNs) is lacking; however, and it remains to be seen whether LINC complex or associated proteins (*e.g.* lamins) bear mechanosensitivity. Stretch-activation could ultimately control motor-protein activity and/or the recruitment of NETs to guide chromatin reorganization using mechanisms similar to focal adhesions^66^.

## METHODS

### Substrate Fabrication

To mimic native and stiffened mechanical environments of adult cardiac tissue^28,29,67^, cell culture dishes were coated with soft or stiff polydimethylsiloxane (PDMS, Sylgard®527, Dow Corning) by using different mixing ratios (base:curing agent): 1:1 (*E*=12.7 ± 5.0 kPa) and 1:4 (*E*=139.7 ± 16.2 kPa). To enable live imaging at high magnification using a 100× objective, PDMS was deposited as thin layer (∼100 µm) in gridded imaging dishes (µ-Dish 35 mm high Grid-500, ibidi). PDMS coated dishes were degassed under vacuum for 30 min, cured for 1h at 80°C, ozone-activated via corona arc-discharge (30s) and coated with reduced growth factor basement membrane matrix (Geltrex®, Gibco) for 1h at 37°C to provide attachment sites similar to the cardiac basement membrane. PDMS stiffness was determined via AFM using a spherical borosilicate glass tip (diameter=10 µm, stiffness=0.85 N/m), and Young’s modulus *E* was calculated using a Hertz contact model^68^.

### Cardiac Cell Isolation and Culture

B6.Cg-Tg (HIST1H2BB/EGFP) 1Pa/J mice (Stock No: 006069) were obtained from Jackson Laboratory. All animal procedures were performed following Institutional Animal Care and Use Committee approval. Embryonic mice hearts were harvested 18.5 days post conception. Hearts were minced and incubated in a digestive mix for 30 min at 37°C. Digestive mix contained 2 mg/ml Papain (P4762, Sigma), 500 µg/ml Liberase TM (05401119001, Roche), 5 mM L-Cysteine (C6852, Sigma) and 10 µg/ml DNase-I (D4263, Sigma). Cardiac cells were isolated through gentle trituration using a 1 ml pipette and cells were cultured on prepared substrates in DMEM-F12 Advanced (Gibco) containing 10% fetal bovine serum (Gibco), 1% penicillin-streptomycin (Gibco) and 25 mM HEPES (Gibco) at a density of 50,000 cells/cm^2^. Samples were incubated at 37°C and 5% CO_2_^69^.

### RNAseq Analysis

Total RNA was extracted from cardiac cultures after 4 days on either soft or stiff PDMS (n=4) using the AurumTM Total RNA Mini Kit (Bio-Rad Laboratories). Libraries were constructed by the Purdue Genomics Core Facilities according to standard protocols using TruSeq Stranded mRNA Library Prep Kit (Illumina). Samples were run on a HiSeq 2500 (Illumina) with 200 bp paired-end reads. The filtered Illumina reads were pre-processed and mapped by the Purdue Bioinformatics Core. Sequence quality was assessed using FastQC (v0.11.2) for all samples and quality trimming was done using FASTX toolkit (v0.0.13.2) to remove bases with less than Phred33 score of 30 and resulting reads of at least 50 bases were retained. The quality trimmed reads were mapped against the bowtie2-indexed reference genome downloaded from Ensembl using Tophat (v2.0.14) with default parameters. RNAseq data can be obtained from the GEO database (GSE109405).

Histone subfamily classifications were done using HistoneDB 2.0 (https://www.ncbi.nlm.nih.gov/research/HistoneDB2.0/). Pathways with enriched differential gene expression (*p*<0.2, FPKM>1) were screened using the KEGG database via the functional annotation tool DAVID (https://david.ncifcrf.gov/). Analyses of functionally grouped networks was performed using the ClueGO (v2.5.0) app on the Cytoscape (v3.6.0) software tool. Genes associated with the GO term *histone modification* (GO:0016570) were obtained from AmiGO 2 (http://amigo.geneontology.org/amigo) and differentially expressed genes (p<0.2) in this ontology were used to screen for child GO terms using AmiGO’s term enrichment tool (v1.8).

### RT-qPCR Analysis

Total RNA was extracted from cardiac cultures after 4 days on either soft or stiff PDMS (n=4) using the AurumTM Total RNA Mini Kit. RNA was reverse transcribed into cDNA via iScriptTM Reverse Transcription Supermix and real-time quantitative PCR was performed with SsoAdvancedTM Universal SYBR® Green Supermix in a CFX96 Touch™ thermocycler (all kits and devices from Bio-Rad Laboratories) using 10 ng of cDNA as input. Primers were custom designed using NCBI primer blast, cross-confirmed in Ensembl gene database and synthesized by IdtDNA. Primers span at least one exon-exon junction. Relative expression change was calculated using the ΔΔCt method. All data was normalized to the reference genes *Gapdh* and *ActB* as established in previous heart studies^70,71^. Primer sequences are listed in **Extended Table 4**.

### Immunofluorescence Staining

Cells were fixed in 4% ice-cold PFA for 10min, permeabilized with 1% Triton-X100 in PBS for 15 min and blocked with 10%NGS, 1% BSA in 0.1% PBT (0.1% Tween-20 in PBS) for 60 min. Primary incubation was performed at 4°C overnight in 0.1% PBT containing 1% BSA. Secondary incubation was performed in primary incubation buffer for 45 min (RT) at a dilution of 1:500. Actin was counterstained with Phalloidin conjugated to either Texas Red-X for embryonic tissues and marker co-localization *in vitro* studies (Life Technologies), Alexa Flour 488 for cardiac sections of hypertrophic mice (Life Technologies) or CF405 for all other *in vitro* cultures (Biotium). Primary antibodies: H3K9me3 (ab8898, Abcam, 1:800), H3K27me3 (ab6002, Abcam, 1:200), RNA polymerase II CTD repeat YSPTSPS phospho S2 (ab24758, Abcam, 1:400) and nesprin-1 (ab24742, Abcam, 1:500).

### Tissue Sectioning and Staining

After harvest, tissues were washed in PBS and fixed in 4% PFA at 4°C overnight. Tissue were washed again in PBS, embedded in 6% agarose and sectioned into 100 µm thin slices using a vibratome (VT1000 S, Leica). Tissue sections were further immunostained as described above.

### Analysis of Chromatin Marker Occupancy

Image stacks of immunostained nuclei were recorded on a Nikon A1R confocal microscope using a 60× oil immersion objective. A custom MATLAB code was used to calculate the intensity of each chromatin marker with respect to distance of the nuclear border. Briefly, the H2b-eGFP image stack was used to determine the nuclear border and the nuclear center for each nucleus. For each pixel, the distance to the nuclear center was calculated and histogram-normalized image intensities for each marker channel were collected. Nuclear center distances were normalized to the maximum distance of each corresponding center trajectory resulting in a normalized center distance of 0 for the center and 1 for the nuclear periphery for any nuclear geometry. Normalized nuclear center distances were then binned in 0.01 steps (100 bins total) and marker intensities for pixels in the same bin were averaged for each nucleus. MATLAB code is available from the corresponding author upon request.

### Hypertrophic Animal Model

Female, 12 weeks old C57BL/6J mice (Jackson Laboratory) were surgically implanted with a mini-osmotic pump (ALZET Model 2004, DURECT Corporation) to deliver either saline (n=5) or AngII (n=4) at a rate of 0.28 µL/h for 28 days. For the AngII group, the AngII powder was dissolved in saline to provide a 1000 ng/kg/min infusion rate. Mice were monitored at baseline and again on day 28 post-surgery using a high resolution, small animal ultrasound imaging system (Vevo2100 Imaging System, FUJIFILM VisualSonics) to assess cardiac size and function^72^. After 28 days, mice were euthanized, hearts harvested, and the left ventricles were cut along the transverse plane. Tissue sectioning and staining was performed as described above. All animal procedures were approved by the Institutional Animal Care and Use Committee.

### Small Animal Ultrasound Imaging

A high resolution, small animal *in vivo* ultrasound imaging system was used to assess cardiac size and function at baseline and again on day 28 post-surgery. Briefly, anesthetized mice (1-3% isoflurane) were positioned supine on a heated stage. Body temperature was monitored using a rectal probe, while cardiac and respiratory rates were monitored using stage electrodes. Hair on the thoracic region was removed with depilatory cream and ultrasound gel was applied. A linear array transducer (MS550D) was used to view both the long and short axes of the heart. Two-dimensional cine loops in brightness mode (B-mode) and motion mode (M-mode) were collected. In addition, high temporal resolution cine loops were acquired using ECG-gated Kilohertz Visualization (EKV). These images were analyzed with Vevo2100 software (FUJIFILM VisualSonics) to determine ejection fraction and fractional shortening.

### Adenovirus Generation and Transduction

Adenoviral vectors for LINC disruption experiments were generated as described before^73^ using the AdEasy vector system. The decoupling and control construct were assembled via Gibson cloning using NEBuilder® HiFi DNA Assembly Master Mix (New England Biolabs). The decoupling construct contained the C-terminal end of the nesprin-3 gene (*Syne3*) including the transmembrane (Tm) and the KASH domain. The N-terminus was replaced by a far-red fluorescence protein (mNeptune2.5) for visualization and a signal peptide (SP, from *Tor1a*) for membrane integration. The control construct was identical to the decoupling construct but lacked the KASH domain. AdEasy plasmids were a gift from Leslie Leinwand. pcDNA3-mNeptune2.5 and pcDNA3-mRuby2 were a gift from Michael Lin (Addgene, #51310 and #40260)^74^. Embryonic cardiac cells were infected at day one with 100×MOI, as determined by plaque assay, and further analyzed at day two or day four as described above. Plasmid maps and stab cultures of adenoviral transfer plasmids can be obtained from Addgene (CTL: #122243 and K3: #122242).

### Nuclear Bulk Linear Strain and Translation

CMs cultured on soft PDMS, stiff PDMS or TCP or infected with K3 or CTL on day one on soft PDMS and image series of nuclei (6.4 fps) were recorded during contraction on day two or four using an inverted epi-fluorescence microscope (Nikon Ti-Eclipse) with a 100× oil immersion objective (0.16 µm/pix) and an iXon^EM^+ EMCCD camera (Andor). A custom MATLAB code was written that tracked nuclear outlines during image series. Bulk linear strain was calculated as the average of the differential length of major and minor axis while translation was determined via centroid tracking. For each nucleus, the results of four contraction cycles were averaged for one data point. MATLAB code is available from the corresponding author upon request.

### Sarcomere Shortening Analysis

CMs cultured on soft or stiff PDMS were infected with K3 or CTL on day one and image series (6.4 fps) of myofibril contractions were recorded on day four using an inverted epi-fluorescence microscope (Nikon Ti-Eclipse) with a 100× oil immersion objective (0.16 µm/pix) and an EMCCD camera (iXon^EM^+, Andor) via expression of α-actinin 2-mRuby2. A custom written MATLAB code was used to quantify the length of different sarcomere features before and during contraction from α-actinin 2 fluorescence intensity profiles stretching 15-20µm and containing 8-12 α-actinin 2-rich Z-disks (**Fig. 6a**). Overall sarcomere lengths were determined as the distance between neighboring peaks. Data points between peaks were interpolated (cubic) to achieve sub-pixel resolution and the mid-intensity between a maximum (peak) and a minimum (valley) was used to separate the Z-disks from A-bands for each segment (**Fig. 6b**). Values along one intensity profile were averaged and the difference in length before and during contraction for each sarcomere feature was calculated. A total of 25 intensity profiles were analyzed for each substrate. MATLAB code is available from the corresponding author upon request.

### Marker Co-Localization Analysis

Cardiac cultures were immunostained after four days on soft PDMS and image *z*-stacks were acquired on a Nikon A1R confocal microscope using a 60× oil immersion objective. *In vitro*, myofibrils primarily intersected with nuclei on the basal side (**Extended Fig. 6**). A custom MATLAB code was used to calculate marker co-localization scores for basal nuclear z-slices. For that, marker channels were binarized (0 or 1) using accumulated histogram thresholding with cut-off values of 90% for actin and α-actinin 2, 95% for H2b, and 85% for H3K9me3 and H3K27me3 to achieve equal representation of true pixels (=1) amongst channels. A marker co-localization score for any marker pair was calculated as the percentage of overlapping pixels (number of pixels that are true for both channels divided by total number of pixels in the nucleus) divided by the probability that positive pixels overlap by chance (independent probability): co-localization score = p(A∩B) / [p(A) × p(B)] with A {marker channel 1=true} and B {marker channel 2=true}. MATLAB code is available from the corresponding author upon request.

### Spatial Intranuclear Strain vs. Chromatin Marker Analysis

After two days on soft PDMS, image series (7.2 fps) of nuclear deformation during CM contractions were recorded on a Nikon A1R confocal microscope using a 60× oil immersion objective (0.14 µm/pix) via the expression of H2b-eGFP. Nuclear frames during diastolic rest and during peak contraction were used as template and target, respectively, to generate high-resolution intranuclear strain maps via *deformation microscop*y developed previously^31,32^. The location of each cell was recorded through the use of gridded imaging dishes (µ-Dish 35 mm high Grid-500, ibidi). After live imaging, cells were fixed, immunostained for chromatin markers, relocated and image color stacks were recorded again on a confocal microscope. To compensate for changes in nuclear morphology during fixation, intranuclear strain maps and chromatin marker maps were aligned via the registration function of the *deformation microscopy* algorithm using the common H2b channel in both datasets and marker channels were interpolated (cubic) to match strain resolution.

A custom written MATLAB code was used to analyze nuclear strains over chromatin marker intensities. For non-continuous analysis (**Fig. 7c**), marker channels were binarized using accumulative histogram thresholding with a cut-off value of 75% and strain values for true marker pixels were averaged. A composite map was generated by assigning pixels to the marker with the highest normalized intensity value, if multiple channels were true, or labeled as unassigned if no channel was true.

For angular analysis, the angle of each pixel with respect to the nuclear center was obtained with the angle of nuclear translation during contraction set to 0°. Nuclear translation angle was determined as the angle of the trajectory that connected the nuclear centers before (resting) and during CM contraction (**Extended Fig. 8**). Pixels were binned in 1° steps and marker intensities for each channel were averaged for each bin. Areas represent SEM between cells. MATLAB codes are available from the corresponding author upon request.

### Western Blot Analysis

After four days on soft or stiff PDMS substrates, cardiac cells were lysed in tris-triton buffer containing a protease/phosphatase inhibitor mix (Sigma). Protein concentrations were determined using Bradford Assay Kit 1 (Bio-Rad) and 30µg of protein were loaded onto 8% SDS-Page Gels (Thermo Scientific), transferred to PVDF membranes (EMD Millipore) and immunodetected using an ECL substrate kit (Life Technologies) and a PXi imaging system (Syngene). Densitometric quantification was performed using ImageJ (v1.50e). Primary antibodies: p130Cas (13846, Cell Signaling), phospho-p130Cas Tyr410 (4011, Cell Signaling) and Gapdh (5174, Cell Signaling).

### Statistical Analysis

One-way (1W) or two-way (2W) ANOVA with Tukey’s Honestly Significant Difference post hoc test or two-tailed t-test analysis was performed to evaluate statistical significance using JMP Pro12 software (SAS Institute). Displayed error (SD=standard deviation, SEM=standard error of the mean), number of individual data points (n), number of independent experiments (exp., if different from n), hypothetical population mean (HM, for single sample t-test), significances and statistical tests used are indicated in the figure captions. Boxplots are shown with all data points overlaid, a square representing the mean and whiskers indicating the data span under the exclusion of outliers (coefficient=3).

## ACKNOWLEDGEMENTS

We are grateful to Leslie Leinwand and Stephen J. Langer for providing materials and technical assistance for the generation of adenoviruses. We also thank Michael Rafuse for technical assistance regarding tissue sectioning. This work was supported in part by grants to C.P.N.: NIH R01 AR063712, NIH R21 AR066230, and NSF CMMI CAREER 1349735.

## AUTHOR CONTRIBUTIONS

Conceptualization, B.S., S.G., C.J.G., S.C., and C.P.N.; Methodology, B.S., S.G., C.J.G., S.C., and C.P.N.; Software, B.S. and S.G.; Formal Analysis, B.S. and S.G.; Investigation, B.S., A.G.B. and S.E.S.; Writing – Original Draft, B.S.; Writing – Review & Editing, All authors; Funding Acquisition, C.P.N.; Resources, C.J.G. and C.P.N.; Supervision, S.G., S.C., and C.P.N.

## EXTENDED DATA

### EXTENDED TABLES

**Extended Table 1.**
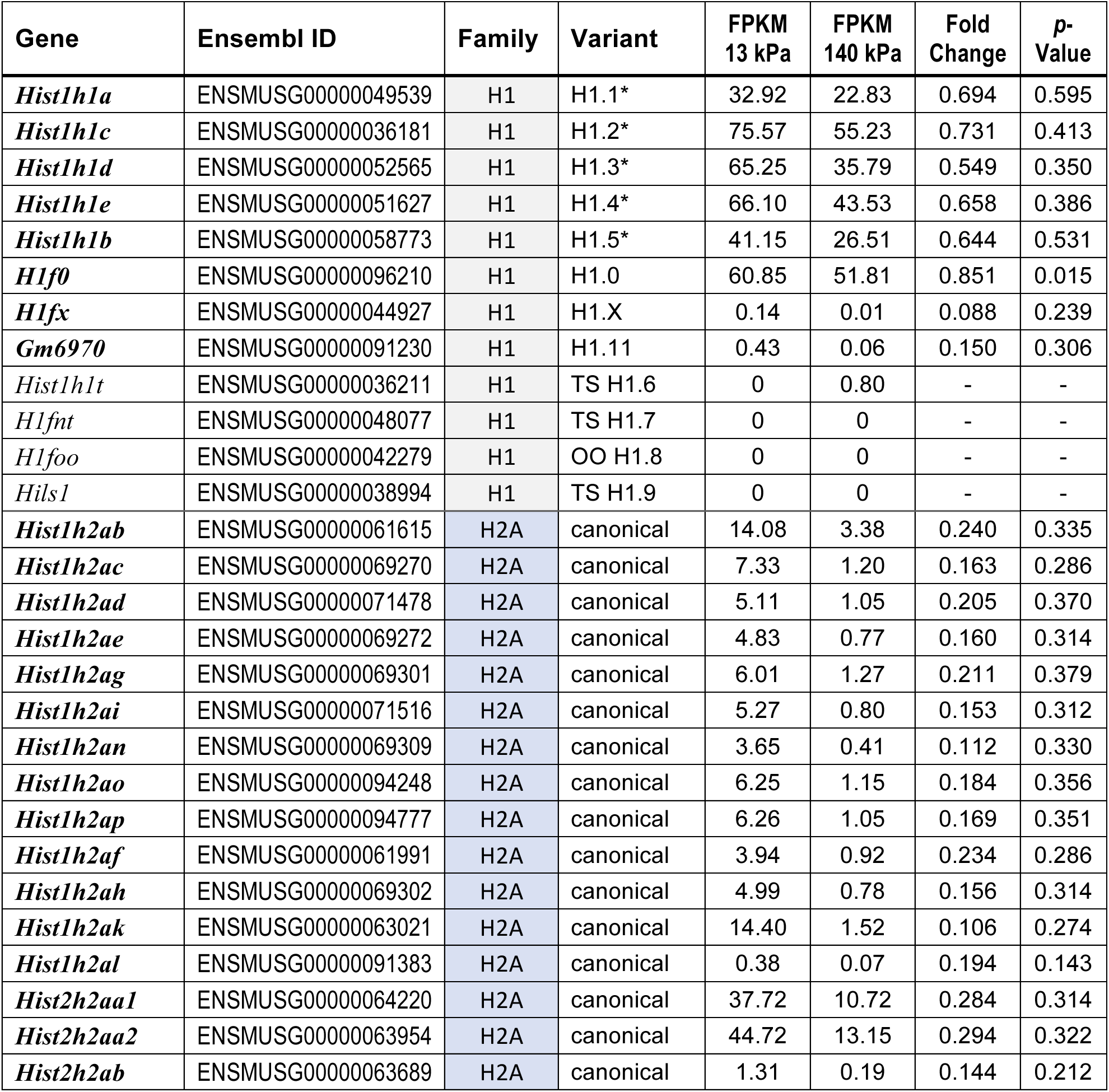
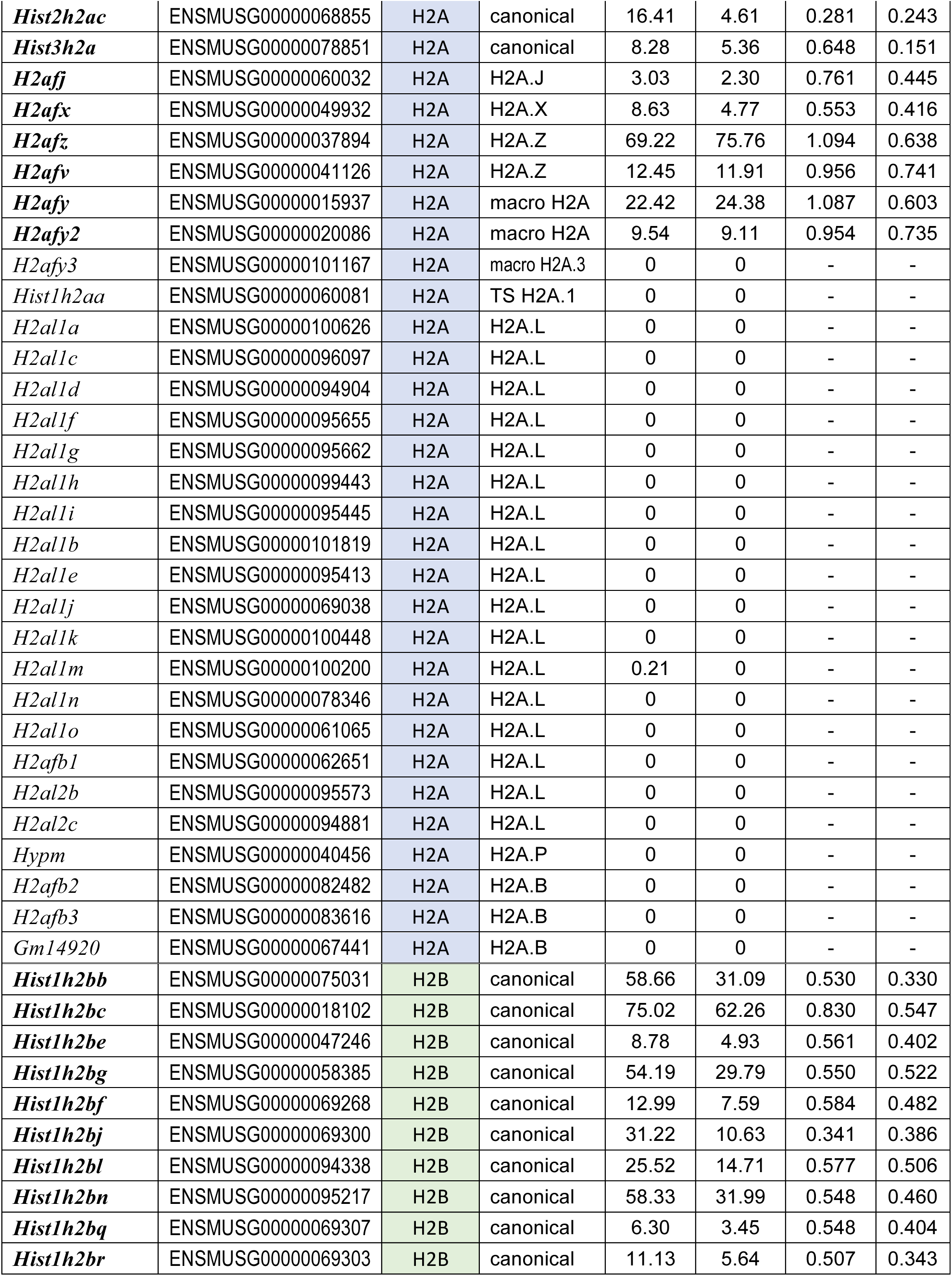
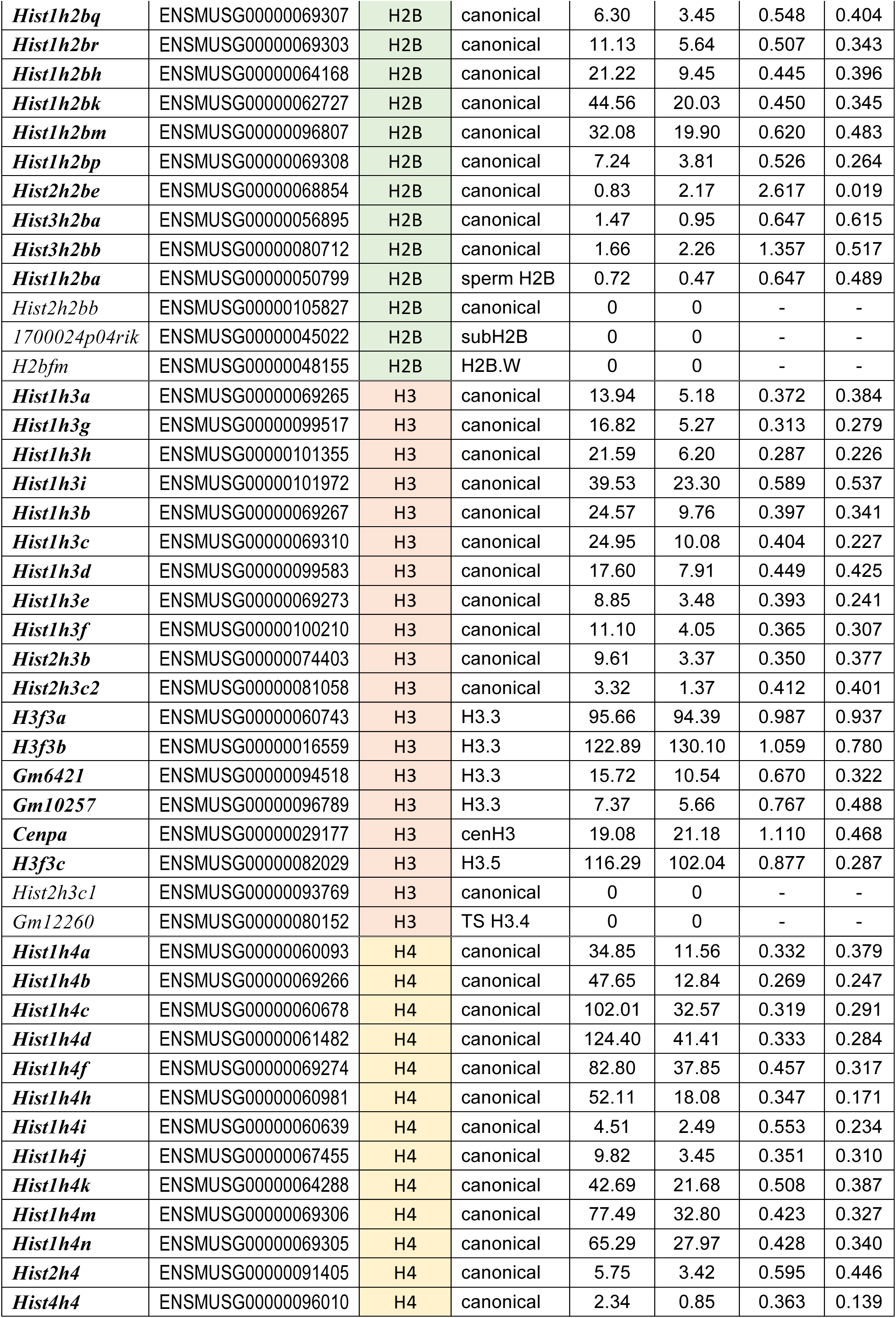
Expression of histone variants in cardiac cultures on cultured on soft or stiff substrates. Embryonic cardiac cells were plated on soft (13 kPa) or stiff (140 kPa) PDMS for four days after which total RNA was harvested to perform RNAseq analysis. Averaged expression (FPKM: Fragments Per Kilobase of transcript per Million mapped reads) for each substrate group, as well as fold changes and *p*-values between substrates, of previously annotated mouse histone variants^36^ are shown; * indicates canonical (generic) H1 variants.

**Extended Table 2.** KEGG pathway analysis of differentially expressed genes from cardiac cultures plated on soft or stiff PDMS substrates. Global gene expression data from RNAseq analysis was used to screen for pathways with enriched differential gene expression (*p*<0.2, FPKM>1) using the KEGG database via DAVID. Listed are pathway terms involved in signal transduction cascades, the number of differentially expressed genes, the percentage of differentially expressed genes compared to the total number of genes associated with that pathway as well as *p*-values and corrected *p*-value Benjamini-scores obtained via the Benjamini-Hochberg procedure.

| **Term** | **Genes#** | **% total** | **$p$-value** | **Benjamini** |
| --- | --- | --- | --- | --- |
| *Signaling pathways regulating pluripotency of stem cells* | 29 | 1.2 | 0.0002 | 0.0650 |
| *Rap1 signaling pathway* | 35 | 1.4 | 0.0050 | 0.2100 |
| *Toll-like receptor signaling pathway* | 20 | 0.8 | 0.0055 | 0.1800 |
| *TNF signaling pathway* | 20 | 0.8 | 0.0120 | 0.3000 |
| *MAPK signaling pathway* | 37 | 1.5 | 0.0210 | 0.2800 |
| *PI3K-Akt signaling pathway* | 48 | 2 | 0.0270 | 0.3200 |
| *Adrenergic signaling in cardiomyocytes* | 24 | 1 | 0.0280 | 0.3100 |
| *HIF-1 signaling pathway* | 18 | 0.7 | 0.0310 | 0.3300 |
| *Cholinergic synapse* | 19 | 0.8 | 0.0340 | 0.3500 |
| *cGMP-PKG signaling pathway* | 26 | 1.1 | 0.0370 | 0.3300 |
| *Insulin resistance* | 18 | 0.7 | 0.0500 | 0.3600 |
| *AMPK signaling pathway* | 20 | 0.8 | 0.0520 | 0.3700 |
| *ErbB signaling pathway* | 15 | 0.6 | 0.0530 | 0.3700 |
| *ECM-receptor interaction* | 15 | 0.6 | 0.0580 | 0.3700 |
| *Neurotrophin signaling pathway* | 19 | 0.8 | 0.0650 | 0.3800 |
| *Estrogen signaling pathway* | 16 | 0.7 | 0.0670 | 0.3800 |
| *Wnt signaling pathway* | 21 | 0.9 | 0.0750 | 0.4000 |
| *VEGF signaling pathway* | 11 | 0.4 | 0.0770 | 0.4000 |
| *Hippo signaling pathway* | 22 | 0.9 | 0.0820 | 0.4100 |
| *Jak-STAT signaling pathway* | 21 | 0.9 | 0.0930 | 0.4300 |
| *Calcium signaling pathway* | 25 | 1 | 0.0960 | 0.4300 |

**Extended Table 3.** Substrate specificity of differentially expressed histone methylating genes. List of differentially expressed (*p*<0.2) genes between soft and stiff PDMS, as determined by RNAseq, with the GO term association *histone methylation* (GO:0016571). Shown are substrate specificities for mono (1), di (2), tri (3) or any methylation of the respective target histone residue as obtained from either UniProt or AmiGO database. H3K9 residues were amongst the most prevalent substrates.

| Gene | Ensembl ID | Substrate specificity | Source | FC | P-value |
| --- | --- | --- | --- | --- | --- |
| *Ehmt1* | ENSMUSG00000036893 | H3K9me1/2 | UniProt | 0.847 | 0.0185 |
| *Paxbp1* | ENSMUSG00000022974 | H3K4me2/3 | UniProt | 0.751 | 0.0666 |
| *Pwp1* | ENSMUSG00000001785 | H3K20me3 | AmiGO | 0.860 | 0.0797 |
| *Setdb1* | ENSMUSG00000015697 | H3K9me3 | UniProt | 0.823 | 0.0830 |
| *Rtf1* | ENSMUSG00000027304 | H3K4me3 | UniProt | 1.127 | 0.1119 |
| *Setd1a* | ENSMUSG00000042308 | H3K4me | UniProt | 0.670 | 0.1190 |
| *Setd1b* | ENSMUSG00000038384 | H3K4me | UniProt | 0.193 | 0.1573 |
| *Eed* | ENSMUSG00000030619 | H3K9me, H3K27me | UniProt | 0.719 | 0.1687 |
| *Arid4b* | ENSMUSG00000039219 | H3K9me3, H3K20me3 | AmiGO | 1.158 | 0.1881 |

**Extended Table 4.**
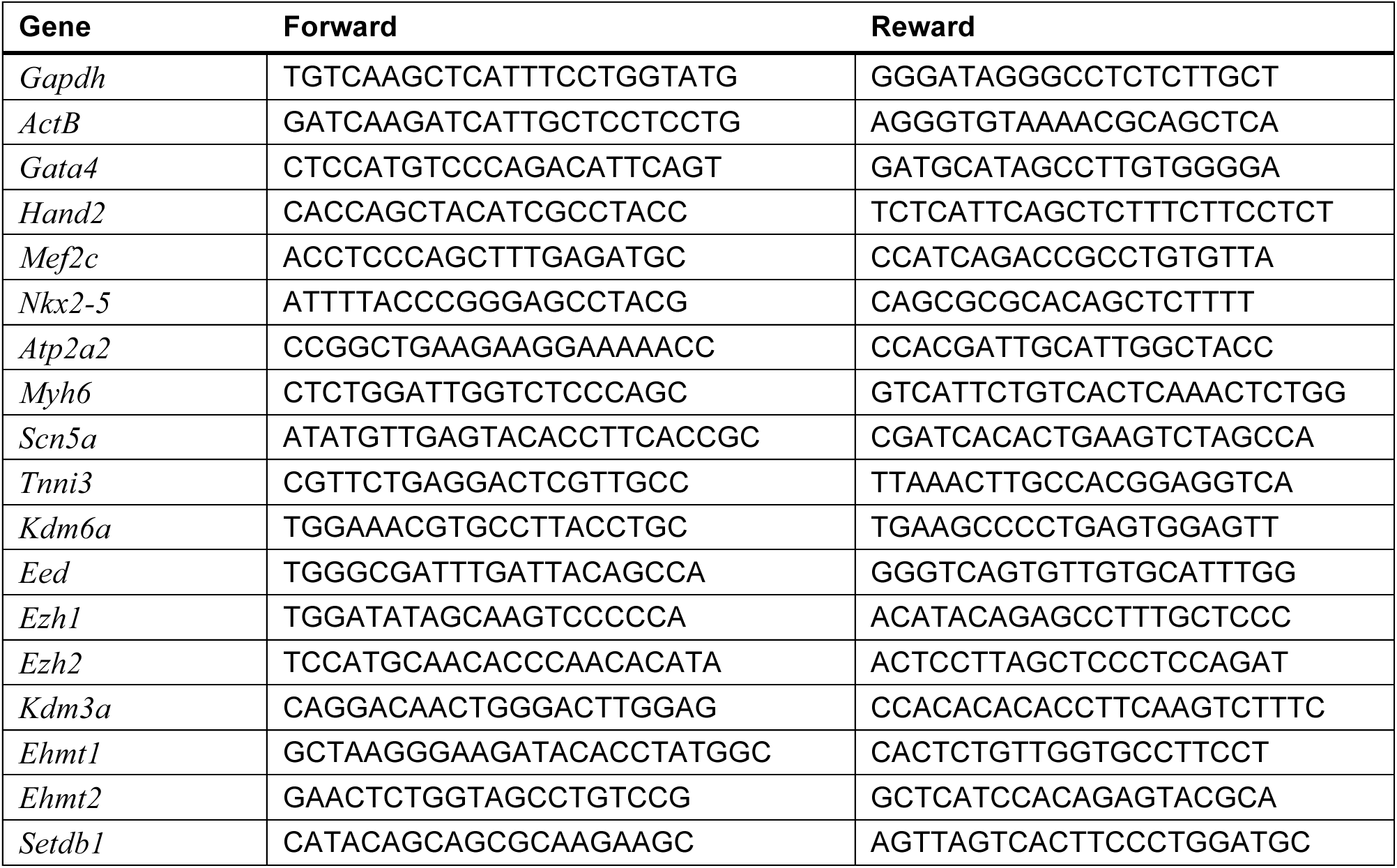
PCR primers for gene expression analysis.

### EXTENDED FIGURES

**Extended Fig. 1.**
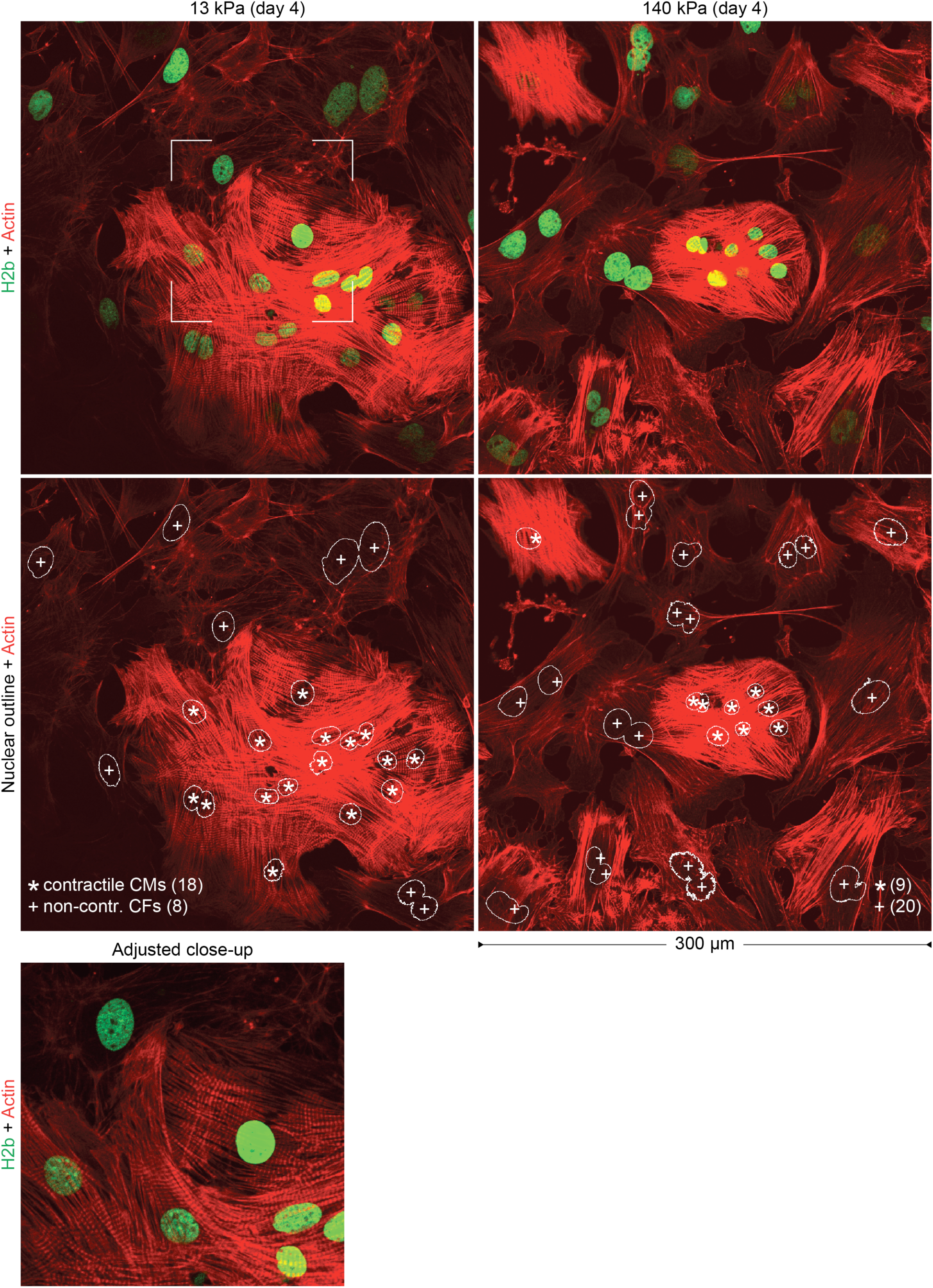
Determining the ratio of contractile CMs to non-contractile CFs in embryonic cardiac cultures on substrates with different stiffness. Embryonic cardiac cells were isolated from (E)18.5 H2b-eGFP embryo hearts and cultured on soft (13 kPa) or stiff (140 kPa) PDMS. After two or four days, cultures were stained for actin and images of 300×300 µm^2^ areas were acquired. Using cell nuclei as reference, cells with clearly formed striated myofibrils were counted as contractile CM (*) or otherwise as non-contractile CF (+). Close-up shows the area indicated by a rectangle in the upper-left frame with adjusted intensity settings to accentuate myofibril striations.

**Extended Fig. 2.**
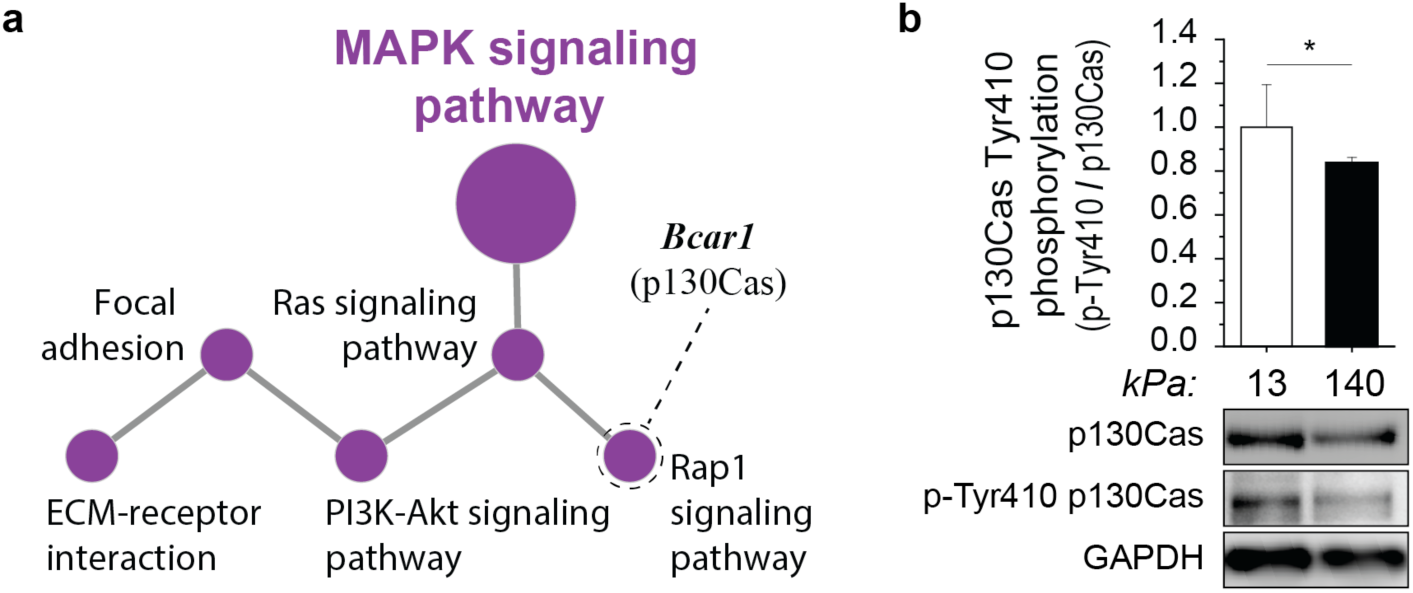
Stiff substrates alter cell-ECM interaction pathways and inhibit stretch-activation of associated p130Cas. Cardiac cells were cultured on soft (13 kPa) or stiff (140 kPa) PDMS for four days before RNA or protein was extracted. **a)** Network analysis of global gene expression change revealed alterations in MAPK signaling and associated pathways that play a role in cell-substrate interaction. Rap1 signaling included the downregulation *Bcar1* coding for p130Cas, a mechanosensitive protein located within the Z-disk lattice in CMs. **b)** Western blot analysis showed reduced tyrosine-410 phosphorylation of the stretch-sensitive mechanosensor p130Cas on stiff PDMS compared to soft; SD; n=3; T-test: * p<0.05.

**Extended Fig. 3.**
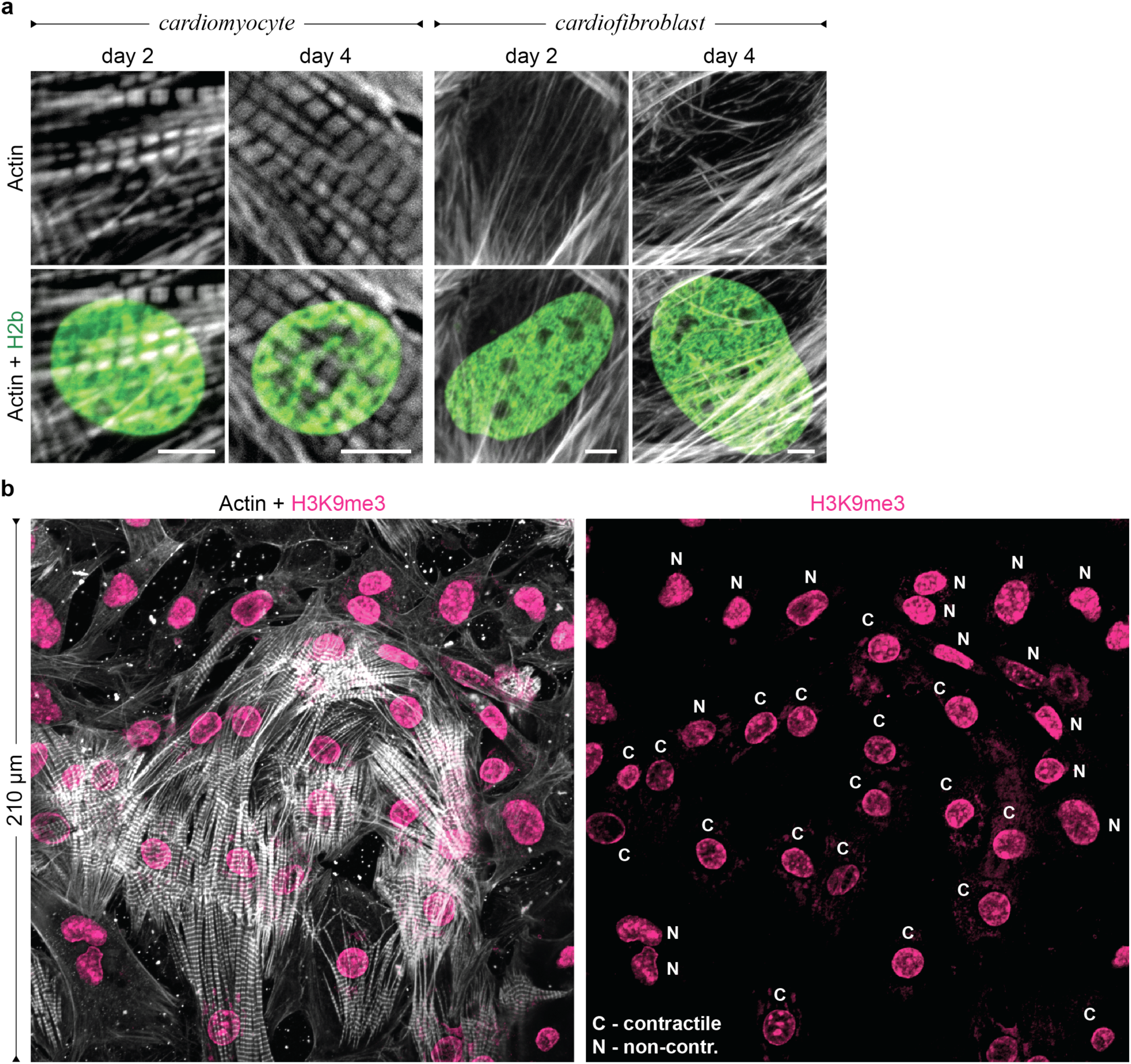
Contractile CMs and non-contractile CFs show differences in the positioning of H3K9 trimethylated chromatin *in vitro*. Embryonic cardiac cells were isolated from (E)18.5 H2b-eGFP embryo hearts and cultured on soft (13 kPa) PDMS substrates. a) Actin staining corresponding to the data from Fig. 3b to distinguish CMs from CFs via the formation of contractile myofibrils; scales=5 µm. b) After two days in culture, contractile (C) CMs with clearly formed myofibrils showed enrichment of H3K9me3-marked chromatin at the nuclear border while actin-fiber forming non-contractile cells (N) showed a more homogenous distribution throughout the nuclear interior.

**Extended Fig. 4.**
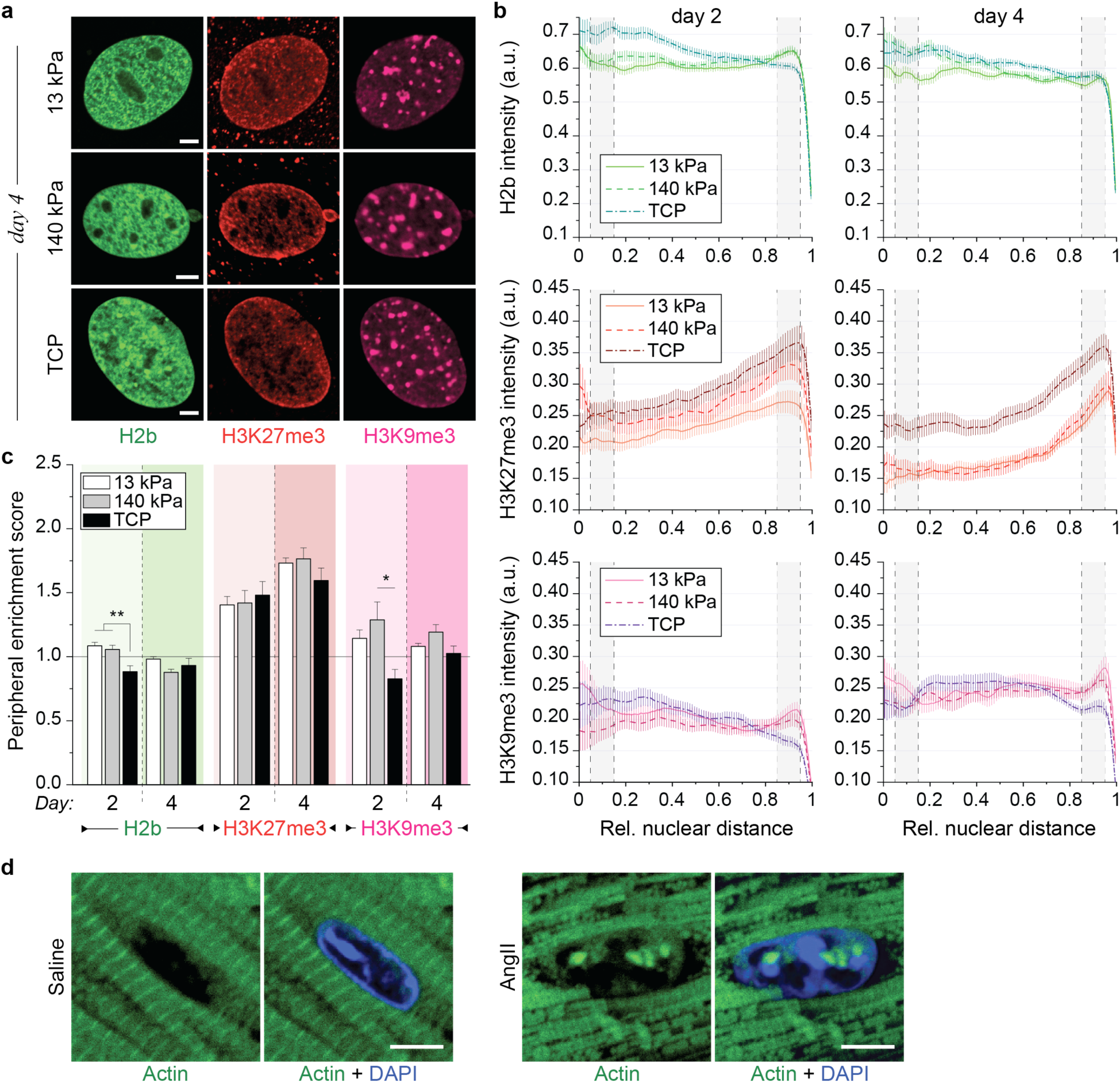
Effect of stiff mechanical environments on chromatin organization in non-contractile CFs *in vitro*. **a)** Embryonic cardiac cells were isolated from (E)18.5 H2b-eGFP embryo hearts and cultured on either soft (13 kPa) PDMS, stiff (140 kPa) PDMS or TCP for two or four days after which cells were stained for H3K27me3 and H3K9me3 as well as actin to distinguish CFs from CMs. **b)** CF nuclei were evaluated for peripheral enrichment of overall chromatin (H2b) or epigenetically marked chromatin using a custom MATLAB code that analyzed marker intensity with respect to its relative distance to the nuclear center (0=center, 1=periphery). Gray areas indicate center and peripheral bin; SEM; n≥30 from 3 exp. **c)** Enrichment scores for each chromatin marker were calculated as the quotient of intensity of the peripheral bin (0.85-0.95) divided by the center bin (0.05-0.15). Substrate stiffness only minorly affected chromatin organization and enrichment of overall and H3K9me3-marked chromatin remained low while enrichment of H3K27me3-modified chromatin remained high throughout the four-day culture period; SEM; n≥30 from 3 exp.; 1W-ANOVA: * p<0.05, ** p<0.01. **d)** Actin staining corresponding to the data from Fig. 4e. All scales=5 µm.

**Extended Fig. 5.**
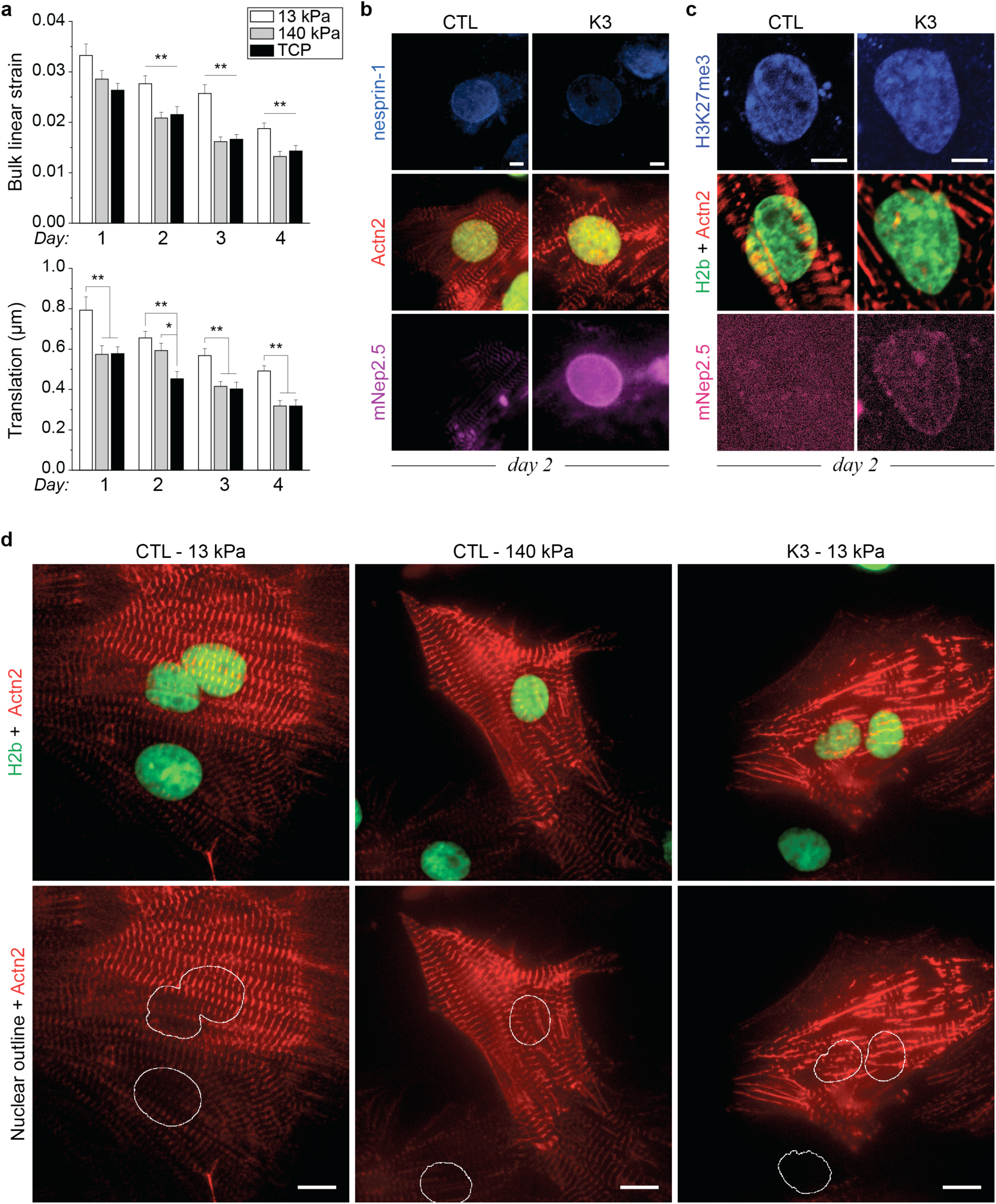
LINC complex disruption in CMs. Embryonic cardiac cells were isolated from (E)18.5 H2b-eGFP embryo hearts. **a)** Image series of nuclei of CMs cultured on either soft (13 kPa) PDMS, stiff (140 kPa) PDMS or TCP for four days were recorded during contraction and bulk linear strain and translational movement of nuclei were determined. Nuclei of CMs cultured on soft substrates showed higher bulk linear strain and translational movement compared to stiff PDMS and TCP; SEM; n>44 from 4 exp.; 1W-ANOVA: * p<0.05, ** p<0.01. **b)** On day one, cells were infected with an adenoviral vector that disrupted LINC connection or a control vector (see Fig. 5a). 24h post infection, widefield images of fixed CMs infected with the decoupling vector showed successful integration of the truncated nesprin construct (mNep2.5) into the outer nuclear membrane while no distinct localization was observed for the control vector. Decoupled cells showed disrupted myofibril formation, particularly around the nucleus, and diminished presence of nesprin-1 at the nuclear membrane; scales=5 µm. **c**) CMs were infected at day 1 and stained for actin and H3K27me3 on day two (shown) and day four. Decoupled cells (K3) showed abolished enrichment of overall and H3K9me3-marked chromatin compared to infected control cells (CTL) while H3K27me3-marked chromatin was similarly enriched (see also Fig. 5); scales=5 µm. **d)** Images of infected cells plated on either soft (13 kPa) or stiff (140 kPa) PDMS. Decoupled cells show disrupted sarcomere fibers, particularly around the nucleus. See also Extended Videos 1-3; scales=10 µm.

**Extended Fig. 6.**
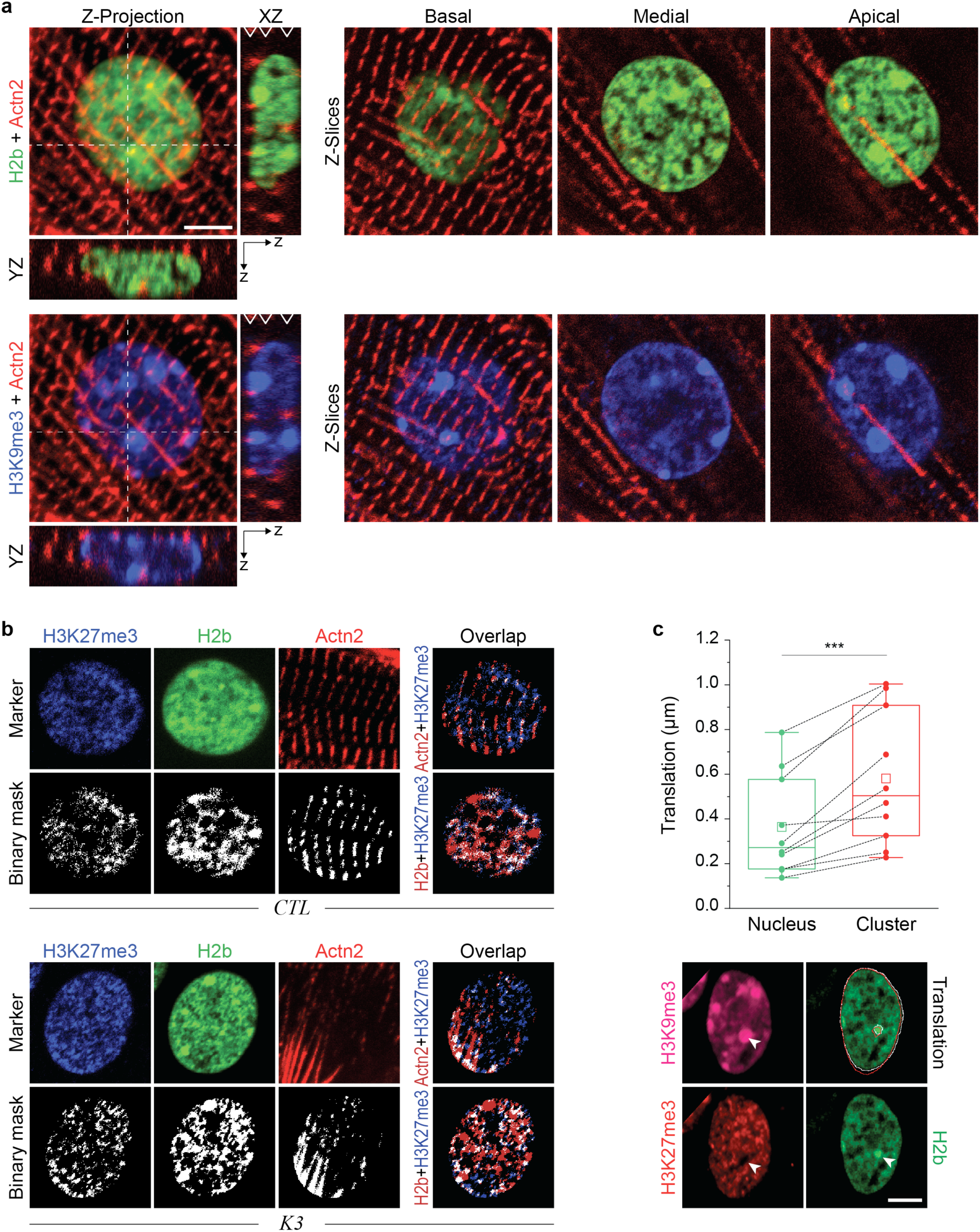
Co-localization of chromatin markers with myofibrils in CMs after LINC complex disruption. Embryonic cardiac cells were isolated from (E)18.5 H2b-eGFP embryo hearts and cultured on soft PDMS (13 kPa). **a)** CMs were infected with K3 or CTL (shown) on day one and stained for H3K9me3 (shown) or H3K27me3 on day four after which z-stacks were recorded on a confocal microscope. Left: Z-projection as well as XZ and YZ slices along white dashed lines for a CTL infected CM. Right: Panels show representative z-slices at different z-positions (basal, medial, apical) indicated by white arrows in the XZ projection. Basal z-slices were used for marker overlap analysis; scale=5 µm. **b)** CMs were infected with K3 or CTL on day one and stained for H3K9me3 (see Fig. 6f) or H3K27me3 (shown) on day four to analyze marker overlap. **c)** Nuclear image stacks of beating CMs were recorded on day two after which cells were stained for H3K9me3 and H3K27me3 and translation of dense, H3K9me3-rich heterochromatin clusters and overall nuclear translation were analyzed. Translational movement was higher for heterochromatin clusters than for nuclei during contractions. White and red outlines indicate nuclear and cluster boundary during rest and peak contraction, respectively; n=10 from 3 exp.; T-test: *** p<0.001; scale=5 µm.

**Extended Fig. 7.**
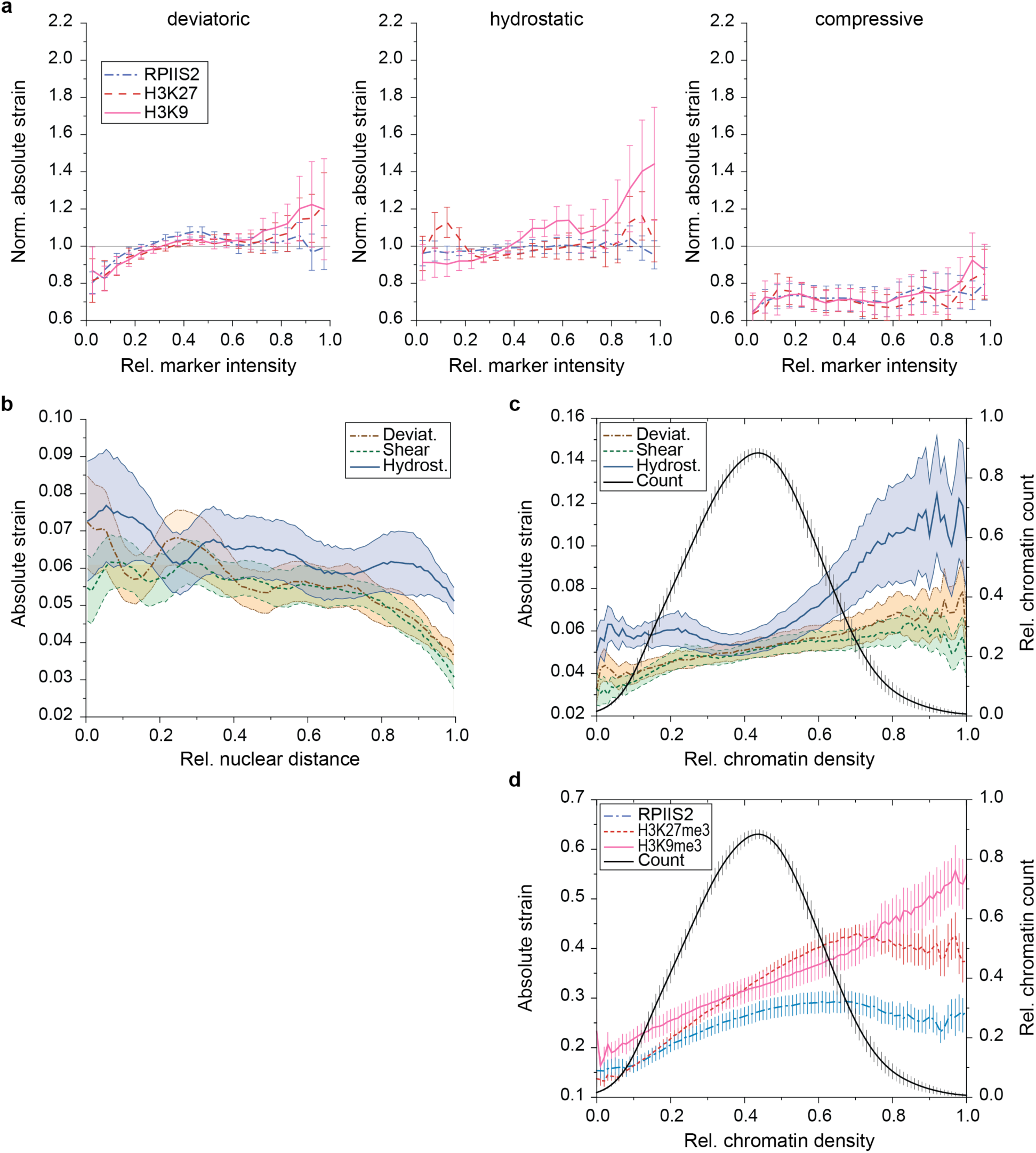
Extended analysis of intranuclear strains during CM contraction. CMs were isolated from (E)18.5 H2b-eGFP embryo hearts and cultured on soft (13 kPa) PDMS for two days. Intranuclear strain maps of CM nuclei during contraction were generated via *deformation microscopy* after which cells were stained for H3K9me3, H3K27me3 or actively transcribed chromatin (RPIIS2) and strain occupancy for chromatin markers was analyzed (see Fig. 7). **a)** Intranuclear strains were analyzed over chromatin marker intensities. **b)** Intranuclear strains were analyzed over distance to the nuclear center. Strains declined towards the nuclear border. **c, d)** Intranuclear strains and chromatin marker intensities were analyzed over relative chromatin density based on H2b intensity. Chromatin density distribution (histogram) is represented as relative chromatin count on the right *y*-axis. Hydrostatic strains are lowest around medium chromatin density (density histogram peak) and increases for denser chromatin which is primarily occupied by H3K9me3 modifications.

**Extended Fig. 8.**
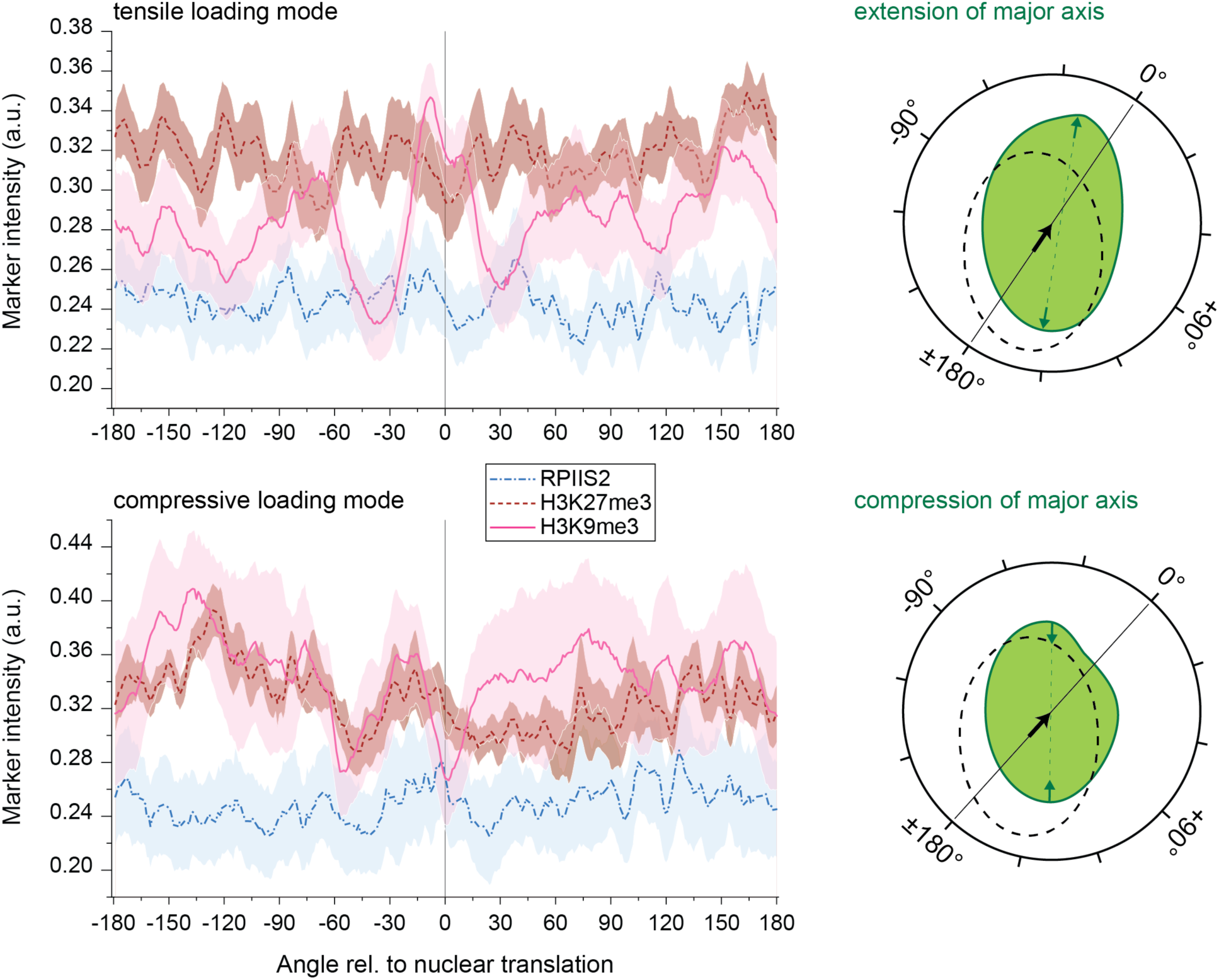
H3K9 methylated chromatin occupancy peaks in the direction of contraction in nuclei with tensile loading mode. CMs were isolated from (E)18.5 H2b-eGFP embryo hearts and cultured on soft (13 kPa) PDMS for two days. Image stacks of CM nuclei were recorded during contractions to determine the direction of nuclear translation. Cells were then stained for chromatin markers H3K9me3, H3K27me3 or actively transcribed chromatin (RPIIS2). Chromatin marker occupancy was calculated with respect to the angle of the nuclear center with the angle of nuclear translation set to 0°. Cells with extended major axis during contraction (tensile loading mode, n=20, same as intranuclear analysis) showed a distinct peak of H3K9me3 intensity ±30° around the direction of translation while a decline in H3K9me3 intensity was observed for cells with shortened major axis (compressive loading mode, n=8). Right side provides a graphic illustration of angular analysis showing nuclear outlines during resting phase (doted black) and peak contraction (solid green). The black arrow indicates the direction of translation, which defines the 0° point, and green arrows demonstrate extension or compression of the nuclear major axis used to determine the loading mode of cells; areas=SEM; from 5 exp.

